# Dynamic trade-offs between biomass accumulation and division determine bacterial cell size and proteome in fluctuating nutrient environments

**DOI:** 10.1101/2022.10.03.510720

**Authors:** Josiah C. Kratz, Shiladitya Banerjee

## Abstract

Bacteria dynamically regulate cell size and growth rate to thrive in changing environments. While much work has been done to characterize bacterial growth physiology and cell size control during steady-state exponential growth, a quantitative understanding of how bacteria dynamically regulate cell size and growth in time-varying nutrient environments is lacking. Here we develop a dynamic coarse-grained proteome sector model which connects growth rate and division control to proteome allocation in time-varying environments in both exponential and stationary phase. In such environments, growth rate and size control is governed by trade-offs between prioritization of biomass accumulation or division, and results in the uncoupling of single-cell growth rate from population growth rate out of steady-state. Specifically, our model predicts that cells transiently prioritize ribosome production, and thus biomass accumulation, over production of division machinery during nutrient upshift, explaining experimentally-observed size control behaviors. Strikingly, our model predicts the opposite behavior during downshift, namely that bacteria temporarily prioritize division over growth, despite needing to upregulate costly division machinery and increasing population size when nutrients are scarce. Importantly, when bacteria are subjected to pulsatile nutrient concentration, we find that cells exhibit a transient memory of the previous metabolic state due to the slow dynamics of proteome reallocation. This phenotypic memory allows for faster adaptation back to previously-seen environments when nutrient fluctuations are short-lived.

## Introduction

In their natural environment, bacteria must be able to sense and adapt rapidly to time-varying environmental stressors to survive and proliferate. Not surprisingly, bacteria exhibit tight regulatory control over their growth physiology and cell morphology [1, 2], and alter both in response to fluctuating nutrient perturbations, resulting in dynamic growth rate and cell size changes in time-varying environments [3–6].

Significant research has gone into understanding how bacterial cell size is coupled to growth rate [7], DNA replication [8, 9], and gene expression [10] at steady-state, and how size homeostasis is maintained despite division and growth rate noise [11, 12]. In addition, characterization of a large portion of the steady-state bacterial proteome across different growth conditions has improved understanding of the resource allocation strategies employed by bacteria in different environments [13–15]. Motivated by experimental data, various coarse-grained models of cell physiology have been developed in recent years, which explain the regulation of cellular growth rate and cell size control from underlying proteome allocation strategies at steady-state [10, 16–19]. However, bacteria do not exist naturally in such conditions, but instead thrive in rapidly changing environments. As a result, it remains unclear how cells sense changes in the environment and dynamically regulate division and growth in response.

Bacteria reallocate their proteome to relieve metabolic or translational bottlenecks and increase growth rate under a given nutrient limitation [20], but do not always allocate resources in order to optimize steady-state growth rate [21]. For example, bacteria maintain a fraction of inactive ribosomes at steady state regardless of nutrient condition, presumably as a reserve which can be deployed to quickly increase growth rate during nutrient upshift [4, 22]. This apparent strategy highlights the challenges of resource allocation in dynamic environments, specifically that organisms must weigh the trade-offs between optimizing growth rate at steady-state and employing mechanisms that are costly at steady-state but that hasten adaptation to environmental changes [4, 23]. In addition, the molecular mechanisms connecting dynamic resource allocation to division control in bacteria are not clear, nor is our understanding of how these allocation strategies are affected by the temporal pattern of environmental fluctuations. Furthermore, it is not clear if bacterial size modulation is simply a byproduct of the complex cellular response to changing environmental conditions, or if it serves as an adaptive mechanism employed by the cell to improve fitness in time-varying environments.

To understand the dynamics of bacterial growth physiology and size control in dynamic nutrient environments, we have developed a coarse-grained proteome sector model which connects gene expression to growth rate and division control, and accurately predicts the cell-level *E. coli* response to nutrient perturbations in both exponential and stationary phase seen in experimental data [5, 24]. This is done by integrating the dynamics of biochemical elements such as amino acids, ribosomes, and metabolic enzymes with decision-making rules for cell division. We applied this model to study how cells allocate intracellular resources dynamically in response to time-varying nutrient conditions, and found that growth rate and cell size control is governed by dynamic trade-offs between biomass accumulation and cell division. Specifically, our model predicts that bacteria temporarily divert resources to ribosome production over division protein production during nutrient upshift, resulting in a temporary delay in division and an overshoot in added cell volume per generation as cells prioritize biomass accumulation. Conversely, in response to nutrient downshift, cells prioritize division over growth, resulting in a rapid decrease in added volume and interdivision time before relaxing to their steady-state values. As a result, population and single-cell growth rates uncouple outside of steady-state, potentially serving as an adaptive mechanism in time-varying environments. Lastly, when simulating pulsatile nutrient conditions, we find that growth rate and cell size recovery time after pulse cessation both increase with increasing pulse duration. Our model suggests that this transient memory of previous environments is a result of the slow dynamics of proteome reallocation, and provides a passive mechanism for faster adaptation in fluctuating environments.

## Results

### Dynamic proteome allocation in time-varying nutrient environment

Bacterial cells integrate time-varying environmental information through a complex set of regulatory networks to control gene expression. Despite this complexity, steady-state proteomics reveals that the expression of proteins with similar function are regulated reciprocally in response to growth rate perturbations, such that various proteome sectors can be defined which coarse-grain the cellular milieu into a limited number of collective state variables [13–15]. To investigate *E. coli* cell size and growth rate control in time-varying nutrient environments, we developed a dynamic model which coarse-grains the proteome into four sectors: ribosomal, metabolic, division, and “housekeeping” (Figure 1A). In this framework, the environment contains time-varying nutrients with concentration *c*, which the cell imports and converts into amino acids using metabolic proteins with protein mass fraction *ϕ_P_*. We assume that the kinetics of protein translation are limited by the abundance of multiple metabolites, and so coarse-grain amino acid abundance into a single group of amino acid precursors, with mass fraction *a*. These precursors are consumed by translating ribosomes, with mass fraction *ϕ_R_*, to synthesize each of the four proteome sectors. As a result, the ribosome mass fraction sets the cellular exponential growth rate, *κ* = dln*M*/d*t* = dln*V*/d*t*, such that growth rate is defined as

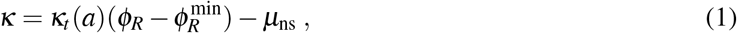

where 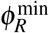 denotes the fraction of ribosomes which are not actively engaged in translation, and *κ_t_* (*a*) is the translational efficiency, which is dependent on amino acid availability such that translation becomes significantly attenuated at low intracellular amino acid levels (see Supporting Information Section I). Here we also coarse-grain the effects of protein turnover and assume that it is governed by a constant, nonspecific degradation rate constant, *μ*_ns_.

**Figure 1.**
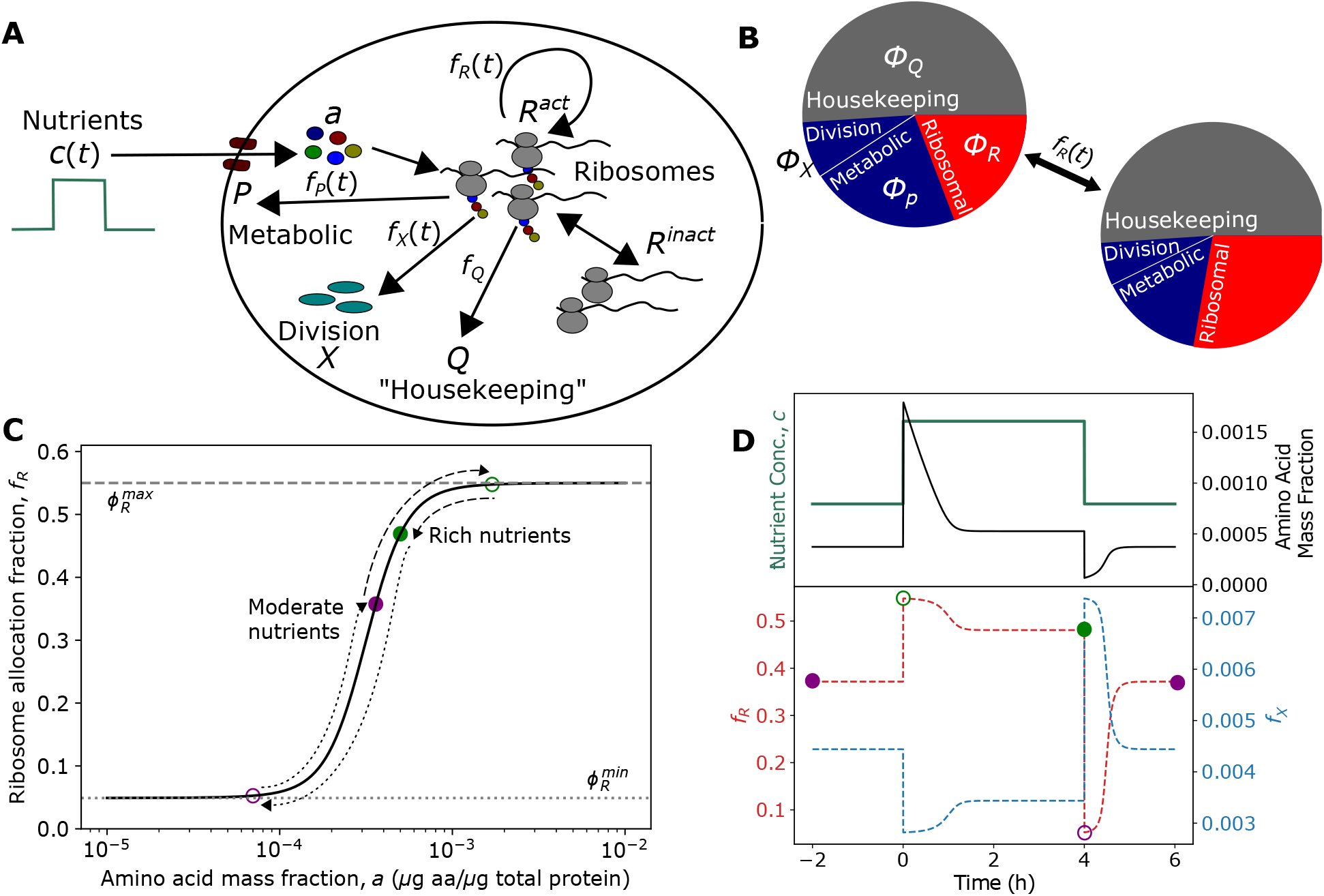
Dynamic resource allocation model for cell growth and division control in dynamic environments. (A) Schematic of coarse-grained model of bacterial cell size control and growth physiology. Nutrients (c) are imported by metabolic proteins (P) and converted to amino acid precursors (a), which are then consumed by ribosomes (R) to produce proteins via translation. Division occurs once a threshold amount of division proteins (X) have been accumulated. (B) By dynamically regulating the fraction of the total translational flux devoted to each proteome sector i, *f_i_*(*a*(*t*)), in response to changes in *a* triggered by environmental changes, the cell alters its proteome composition, and thus its size and growth rate. (C) The dependence of *f_R_* on *a* is the given by their steady-state relationship. The path of *f_R_* in response to a nutrient-rich pulse is shown with colored circles corresponding to the timepoints shown in (D). *f_R_* is initially given by its steady-state value in poor media (purple, closed). A shift to rich media results in a transient increase in *f_R_* close to its maximum value (green, open), before relaxing back to its new steady-state value (green, closed). The path during upshift is given by the dashed line. A shift back to poor media results in a temporary drop in *f_R_* close to its minimum value (purple, open), before relaxing back to the original steady-state value (purple, closed). The path during downshift is given by the dotted line. (D) Representative dynamics of amino acid mass fraction (top) and proteome allocation fractions (bottom) during a nutrient pulse. See Table I for a list of parameters. *f_X_* is calculated by assuming that division timing is set by the protein FtsZ (see Supporting Information section V).

In response to changes in nutrient availability, the cell reallocates its proteome by altering the fraction of translational flux, 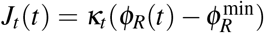, devoted to each sector, such that the time dynamics of each sector can be written as

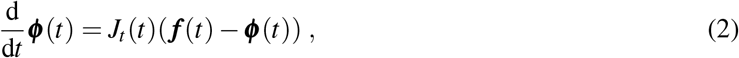

where the vectors ***ϕ***(*t*) = [*ϕ_R_*(*t*), *ϕ_p_*(*t*), *ϕ_X_*(*t*),*ϕ_Q_*(*t*)] and *f*(*t*) = [*f_R_*(*t*), *f_P_*(*t*), *f_X_*(*t*), *f_Q_*(*t*)] denote the protein mass fraction and translational flux allocation fraction of each sector at time *t*, respectively. Proteomics data from *E. coli* reveal that a significant fraction of the proteome is invariant to environmental perturbations [13]. As a result, we define the “housekeeping” sector such that it contains all the proteins whose proteome allocation is not growth rate dependent. Consequently, the mass fraction, *ϕ_Q_*, and allocation fraction, *f_Q_*, are equal and remain constant. This assertion constrains the dynamics of flux allocation such that *f_R_*(*t*) + *f_P_*(*t*) + *f_X_*(*t*) = 1 – *f_Q_* = *ϕ^max^*.

To model division control, we employ a threshold accumulation model of cell division in which division is triggered after a cell accumulates a fixed number of division proteins (collectively referred to as X proteins) [10, 17, 25, 26]. Since the total protein abundance per cell scales with growth rate [7, 8] and if the threshold remains constant [5], the average protein mass fraction of division proteins per cell necessarily decreases to maintain the constancy of the threshold, and thus must be part of the metabolic sector [12, 17]. Consequently, we assume that allocation to the division sector, *f_X_*(*t*), is given by a linear combination of a basal allocation fraction, *β*, and a time-dependent fraction, 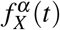, whose expression is co-regulated as part of a larger metabolic sector, 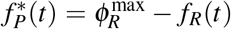. As a result, the flux allocation constraint can be simplified such that 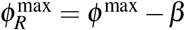, where 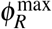 represents the upper limit to allocation fraction devoted to ribosomal proteins. Using the simplified constraint, *f_X_*(*t*) can be expressed such that its time dependence is solely through *f_R_*(*t*), yielding

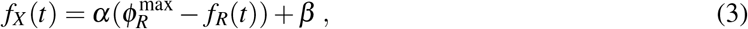

where *α* is the fraction of the co-regulated sector 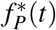 made up of division proteins. From Eq. (3), we see that when the fraction of cellular resources allocated to ribosome production increases, metabolic and division protein translational flux is necessarily downregulated, and vice versa (Figure 1B). This highlights the trade-off that cells must make between biomass accumulation and division in dynamic environments.

Critically, as all other proteome sectors are defined in terms of *f_R_*(*t*), the time-dependence of *f_R_* must be specified. To do so, we assume that dynamic reallocation is driven by gene-regulatory networks which are dependent on the amino acid pool, such that the time dependence of *f_R_* is given through its dependence on the time-varying amino acid mass fraction *a*, thus *f_R_*(*t*) = *f_R_*(*a*(*t*)). The dynamics of *a* are coupled to Eq. (2) and are given by the difference in the metabolic and translational fluxes, such that d*a*/d*t* = *J_n_ – J_t_*, where the metabolic flux, *J_n_*, is proportional to the metabolic sector mass fraction, *ϕ_P_*. Using the proteome constraint relationship above, the dynamics of *a* can be written in terms of the proteome mass fractions, such that

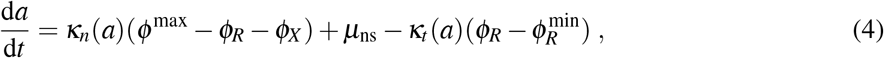

where *κ_n_*(*a*) is the nutritional efficiency (see Supporting Information Section I), and is dependent on *a* such that nutrient import becomes significantly attenuated at high values of *a* to reflect end-product inhibition of biosynthesis pathways and inactivation of nutrient importers at high intracellular amino acid concentrations [27].

Changes in environmental nutrient availability result in a flux imbalance which alters the size of the amino acid pool. In this way, *a* acts as a read-out of flux imbalance, and so by altering proteome allocation in response to *a*, the cell can dynamically respond to nutrient changes. To obtain the functional form of *f_R_*(*a*(*t*)), we assume that *a*(*t*) sets the allocation fraction *f_R_*(*a*(*t*)) via the steady-state relation 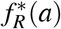, such that 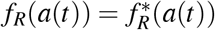. Furthermore, we assume that the cell maximizes translational flux at steady-state, which allows us to express *f_R_* solely in terms of the amino acid mass fraction, *a*. 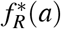 is shown graphically in Figure 1C, and predicts that proteome allocation is altered to reduce growth bottlenecks. Namely, when *a* is high, ribosome synthesis is prioritized in order to increase translation flux, but when *a* is low, metabolic protein synthesis is prioritized to increase nutrient import. Mathematically, any monotonically increasing function for *f_R_*(*a*) will produce this type of regulatory behavior. However, by choosing *f_R_*(*a*) to be given by 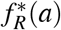, we also ensure that translational flux is maximized at steady-state. This assumption of growth-rate maximization at steady-state has proved fruitful in previous theoretical models to explain bacteria growth laws [16, 27–31], and has been observed experimentally to be the case for many nutrient-limiting conditions [21]. Furthermore, it has been experimentally observed that *E. coli* cells evolve their metabolism towards a state that maximizes growth rate [32–34].

Using the above framework, the dynamics of proteome allocation can be simulated in time-varying nutrient environments by numerically solving the coupled Eqs. (2) and (4) (Figure 1D). In response to a pulse of nutrients, allocation to ribosome synthesis increases drastically to its maximum value before slowly relaxing to its steady-state value in rich nutrients (Figure 1D). In contrast, allocation to division protein synthesis drops significantly before slowly relaxing to a lower steady-state value. Following cessation of the nutrient-rich pulse, the opposite trends occur for each allocation fraction, resulting in an overshoot in *f_X_* and undershoot in *f_R_* before both returning to their initial values (Figure 1D). Mechanistically, this regulation of ribosome expression is carried out by the signaling molecule guanosine tetraphosphate (ppGpp). ppGpp is synthesized when charged tRNA levels are low [35, 36]. As charged tRNA abundance is proportional to amino acid levels, ppGpp thus indirectly acts as a sensor of the amino acid pool. As a result, amino acid abundance is inversely proportional to ppGpp concentration, such that [ppGpp] ∝ 1/*a*. In response to decreased amino acid levels, ppGpp levels increase and repress rRNA expression [35, 36]. Free ribosomal proteins, which can no longer bind rRNA, bind to their own mRNA and suppress additional ribosome translation [37]. Conversely, when amino acids are abundant, ppGpp levels decrease which de-represses ribosome production. In this way, the cell is able to regulate gene expression, and thus translational flux, by responding to changes in amino acid concentration.

### Growth-rate dependent increase in cell size arises from trade-off between biomass accumulation and division protein synthesis

To test the validity of our resource allocation model, we first examined if the model can reproduce experimentally observed steady-state physiological behaviors of bacterial cells, in particular the increase in average cell size with growth rate under nutrient perturbations (Figure 2A) [1, 7, 8]. To model cell size control, the dynamics of proteome allocation must be connected to the dynamics of the total number of division proteins per cell, *X*(*t*), as cells divide at *t* = *τ* after accumulating a fixed number of X proteins, *X*(*τ*) = *X*_0_. Using the relation *X* = *ϕ_X_Vρ_c_/m_X_*, where *ρ_c_* is the protein mass density of the cell and *m_X_* is the mass of division molecule X, the dynamics of *ϕ_X_* can be used to identify the dynamics of the fraction of the total number of division proteins required to trigger cell division, 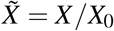,

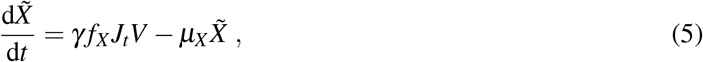

where *γ* = *ρ_c_/X*_0_*m_X_* and *μ_X_* is the degradation rate of the division protein. We thus identify the division protein synthesis rate per unit volume as *k_P_*(*t*) = *γf_X_*(*t*)*J_t_*(*t*). By numerically solving proteome allocation and volume dynamics in conjunction with the division rules given by Eq. (5), single cell size and growth rate dynamics can be simulated in fluctuating nutrient environments.

**Figure 2.**
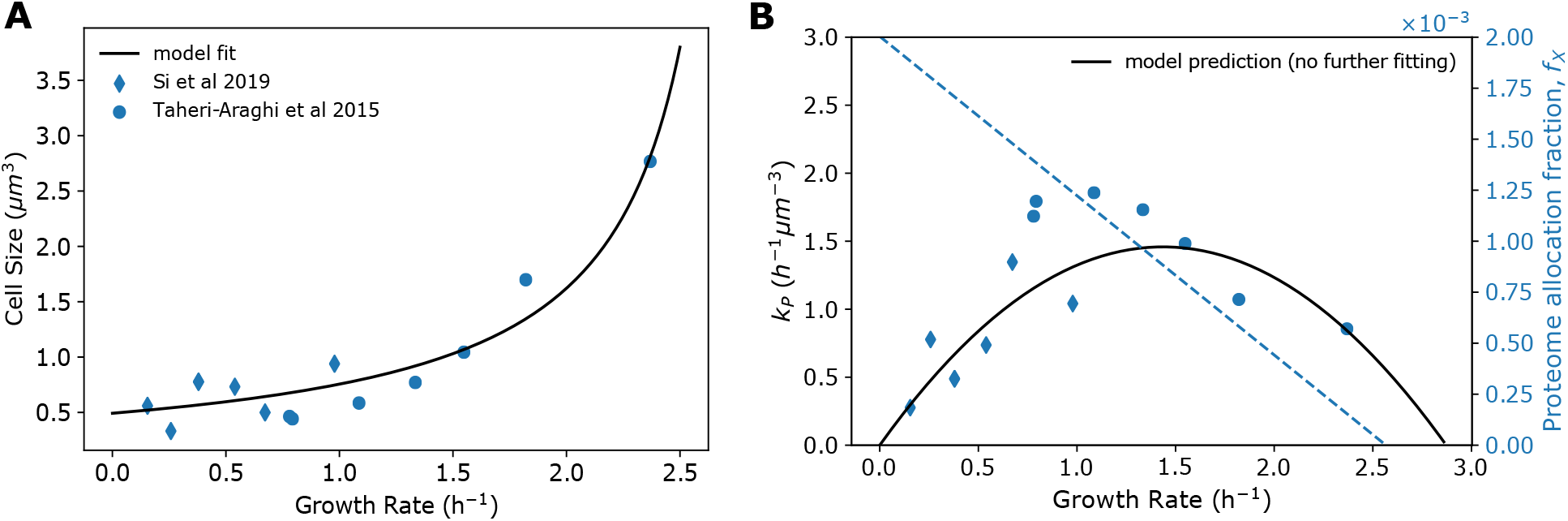
Growth-rate dependent tradeoff between biomass accumulation and division protein synthesis sets steady-state bacterial cell size. (A) Steady-state relationship between population average cell size at birth and growth rate. Solid line shows best fit of Eq. (7), yielding parameters *γα, γβ*, and *κ_t_*, which are given in Table I. Experimental data are taken from Refs. [8, 12]. (B) Non-monotonic dependence of the division protein production rate, *k_P_*, on growth rate, where *k_P_* is estimated from experimental data as 〈*κ*〉/〈*V*_0_〉. Solid line given by Eq. (6), with parameters given by best fit from (A). The allocation fractions to the division protein sector is shown by dotted lines.

At steady-state, *f_R_* = *ϕ_R_* (Eq. 2), allowing the rate of division protein synthesis *k_P_* to be written solely as a function of growth rate. In moderate to fast exponential growth conditions, the effects of protein degradation are negligible. Thus assuming *κ* ≫ *μ_X_* and *κ* ≫ *μ*_ns_, we arrive at

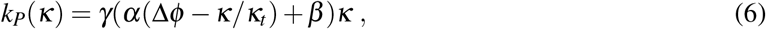

where 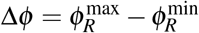. When *κ* ≫ *μ_X_*, this model recapitulates the adder principle employed by *E. coli* to achieve size homeostasis [12], in which a constant amount of volume, Δ*V*, is added each generation irrespective of birth size, Δ*V* ≈ *V*_0_ ≈ *κ/k_P_*. We discuss deviations from this size control behavior in slow growth conditions, when degradation effects become important, in the last Results section. Substituting Eq. (6) into the expression for birth size yields a novel formulation of the size law [7], which links nutrient-limited growth rate to cell size (Figure 2A), such that

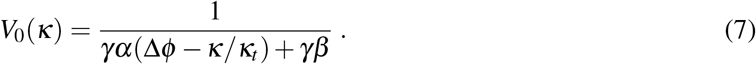

By fitting Eq. (7) to experimental data [8, 12] (Figure 2A), we determine the unknown model parameters *γα, γβ*, and *κ_t_* (see Table I), which allows us to numerically predict the dependence of *k_P_* on *κ*. Interestingly, Eq. (6) predicts a non-monotonic dependence of the division protein production rate on growth rate, which is seen in experimental data when considering a wide range of growth rates (Figure 2B). This behavior can be understood by considering the effects of both *f_X_* and *J_t_*, where here *J_t_* = *κ* when degradation effects are negligible. As growth rate decreases, translational flux allocation to division protein production, *f_X_*, increases while overall translational flux, *J_t_*, decreases (Figure 2B). As such, at fast growth rates, decreasing *κ* results in an increase in *k_P_* due to an increase in *f_X_*. Conversely, at slow growth rates this increase in *f_X_* is dominated by the decrease in *J_t_*, resulting in a reduction in *k_P_*. At intermediate growth rates translational flux and allocation are simultaneously moderately high, consequently yielding the maximum *k_P_* value.

**Table I.** Model parameters. See Supporting Information for more details.

| Parameter | Description | Value | Growth condition | Figure number |
| --- | --- | --- | --- | --- |
| $\phi_R^{\min}$ | inactive ribosome fraction [16] | 0.049 | all | all |
| $\phi_R^{\max}$ | maximum flux allocation to ribosome production [16] | 0.55 | all | all |
| $a_t$ | translation attenuation threshold [27] | $10^{-4}$ | all | all |
| $a_n$ | feedback inhibition threshold [27] | $10^{-3}$ | all | all |
| $\kappa_t^0$ ($\text{h}^{-1}$) | translational efficiency rate constant | 2.6 | all | 1, 3-6 |
|  |  | 5.1 | all | 2 |
| $\kappa_{n,low}^0$ ($\text{h}^{-1}$) | nutritional efficiency rate constant in nutrient-poor media | 5.29 | exponential | 1, 3, 4, 6 |
|  |  | 0 | stationary | 5 |
| $\kappa_{n,high}^0$ ($\text{h}^{-1}$) | nutritional efficiency rate constant in nutrient-rich media | 10 | all | 1 |
|  |  | 62.8 | all | 3-6 |
| $\mu_{ns}$ ($\text{h}^{-1}$) | nonspecific degradation rate | 0 | exponential | 1, 2-4, 6 |
|  |  | 0.1 | stationary | 5 |
| $\mu_X$ ($\text{h}^{-1}$) | division protein degradation rate | 0.6 | all | 1, 3-6 |
| $\gamma\alpha$ ($\mu\text{m}^{-3}$) | fitting parameter, contribution to $k_P$ from co-regulated portion of $f_X$ | 2.3 | all | 1, 3-6 |
|  |  | 3.6 | all | 2 |
| $\gamma\beta$ ($\mu\text{m}^{-3}$) | fitting parameter, contribution to $k_P$ from basal allocation fraction of $f_X$ | 0.7 | all | 1, 3-6 |
|  |  | 0.2 | all | 2 |

The expression for cell volume given in Eq. (7) predicts a maximum growth rate given by *κ*_max_ = *κ_t_*(Δ*ϕ* + *β/α*). This theoretical maximum, however, is nonphysical as it assumes that *f_X_* = *ϕ_X_* = 0, which is never the case (Eq. 3). Growth rate is maximum when 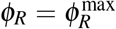, thus giving an upper limit to the physical growth rate at *κ*_max_ = *κ_t_*Δ*ϕ*. Eq. (7) also implies that there is no bound on cell size. However, our expression for *f_X_* constrains cell size to a finite value. When allocation to ribosomes is maximal, *ϕ_X_* = *β*, such that the maximum birth volume *V*_0_ is given by 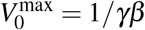.

### Cells transiently prioritize biomass accumulation over division during nutrient upshifts

Recent experimental results show that in response to nutrient upshift, bacteria transiently delay division before increasing to a faster division rate in nutrient-rich media [5]. This behavior is seen clearly in the overshoot in the average interdivision time (*τ*) and added volume (Δ) (Figure 3C-D). We hypothesized that this delay in cell division upon nutrient upshift results from cells prioritizing ribosome production over production of division proteins and metabolic proteins. Using our four-component proteome sector model, we simulated single-cell growth and size dynamics in response to nutrient upshift, and were able to quantitatively capture the experimental results (Figure 3A), as well as predict the dynamics of flux allocation and proteome composition. Importantly, our model was also able to capture growth rate dynamics during both upshift and downshift in many other experimental conditions examined recently by Erickson et al. [18] (Supplementary Figure 3).

**Figure 3.**
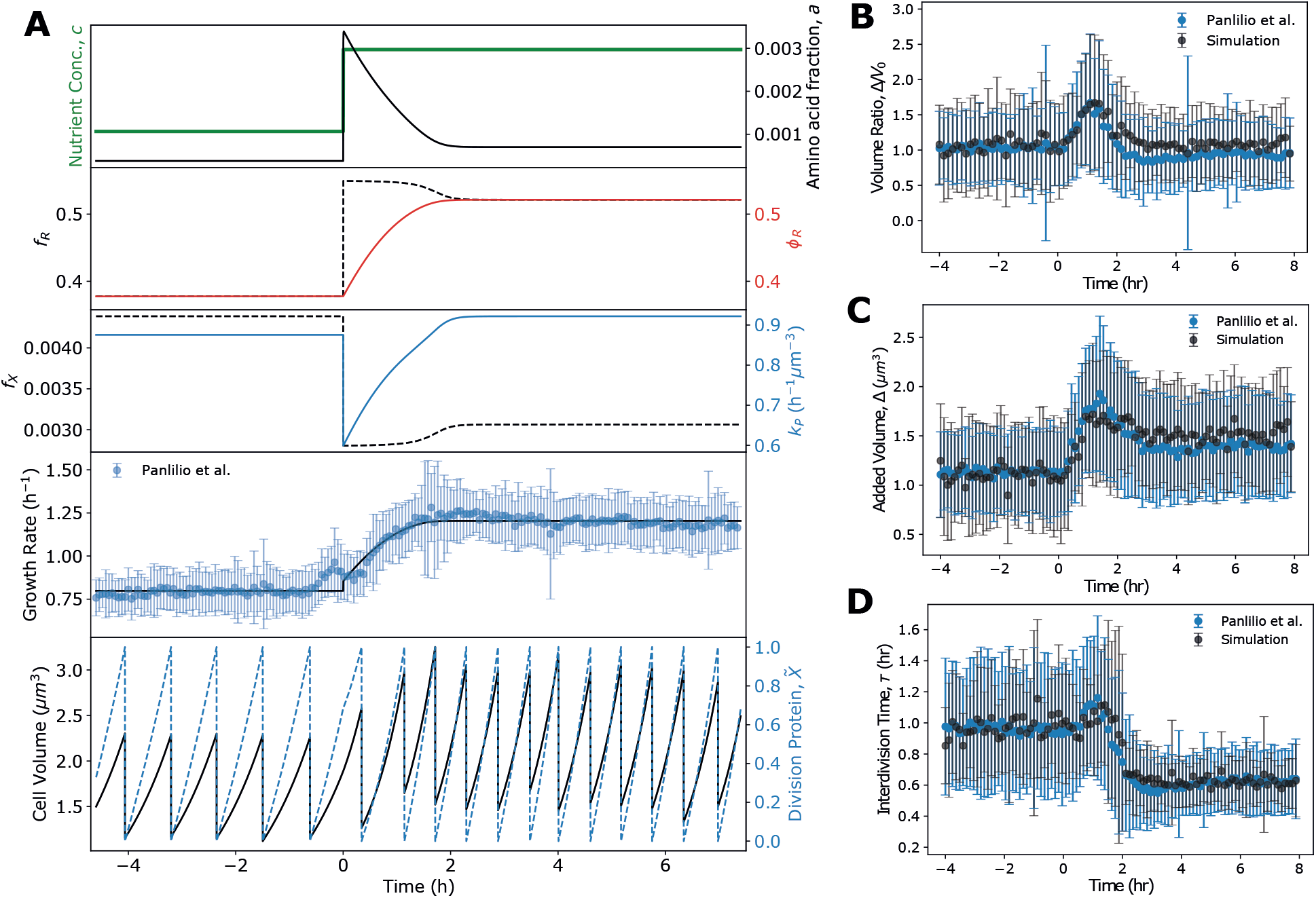
Cell size and division dynamics during nutrient upshift. (A) Simulation dynamics of average amino acid mass fraction, proteome composition, allocation fraction, and division protein production for *E. coli* cells experiencing nutrient upshift. Model parameters obtained by fitting growth rate dynamics to experimental data [5], and are provided in Table I. Bottom: Single-cell volume trajectories were simulated using the model by implementing division rules appropriate for *E. coli*. (B-D) Generation-averaged dynamics of cell volume ratio (B), added volume (C), and interdivision time (D) from single-cell volume trajectories agree well with experimental data. Error bars indicate the standard deviation of the time-binned mean for all time series.

We simulated stochastic single-cell volume trajectories by introducing both growth rate and division noise during the cell cycle, in which only one daughter cell was tracked after each division event (Figure 3A, bottom panel; see Supporting Information Section III). The single-cell level simulations quantitatively capture the average value and noise profile of added volume (Δ), volume ratio (Δ/*V*_0_), and the interdivision time (*τ*) dynamics seen experimentally (Figure 3B-D). In particular, the simulations reproduce the over-shoot in added volume and interdivision time following the nutrient upshift. As hypothesized, our model predicts that in response to increased nutrient availability, bacteria transiently divert resources away from division and metabolic protein production and instead prioritize ribosome production. This regulatory behavior occurs because an increase in nutrient availability transiently causes a mismatch in the translational and metabolic fluxes, yielding a significant increase in the size of the amino acid pool, *a*. In response to the increase in *a*, the cell allocates translational flux to ribosome production at the expense of division and metabolic protein production (Figure 1C-D). This is seen in the temporary drop in division protein production rate, *k_P_*, and overshoot in ribosomal flux allocation, *f_R_*, during the time period during which growth rate increases, before both *k_p_* and *f_R_* relax to their new steady-state values (Figure 3A). Consequently, during this transitional period, bacteria delay division and add significantly more biomass than their birth size (Figure 3B). Importantly, a model in which division protein allocation is constant could not reproduce the observed experimental results, and instead predicted that cell size is invariant to nutrient perturbations (Supplementary Figure 4).

### Growth-rate and cell size recovery time increases with nutrient pulse duration

In order to predict bacterial growth rate and cell size control in more complex time-varying environments, we simulated single-cell trajectories experiencing a pulse of nutrient-rich media with duration *τ*_feast_. For each trajectory with pulse-length *τ*_feast_, we measured the time required (*τ*_recovery_) for both the growth rate and cell volume added per generation to return to the pre-shift level following downshift (Figure 4A,B). Interestingly, in both cases *τ*_recovery_ increased with increasing *τ*_feast_ until saturating at a constant value (Figure 4D), showing that bacteria transiently retain memory of the previous nutrient environment across generations, allowing for quicker recovery to optimal steady-state growth when experiencing short timescale perturbations in nutrient quality. As cellular growth rate is determined by ribosome abundance (Eq. 1), we hypothesized that this phenotypic memory is conferred by the slow dynamics of proteome reallocation and thus ribosome accumulation, which occur on a significantly slower timescale than translational flux reallocation due to the half-life of proteins far exceeding that of mRNA [38]. As such, even though translation proceeds largely at the same rate as transcription in bacteria [39], the stability of previously translated genes allows for transmission of previous metabolic information across time by increasing the time required to reshape proteome composition [23]. In agreement with this hypothesis, we found that the time period over which cells maintain a memory of the previous state is equal to the time required to reshape the proteome to become optimal in the new environment (Figures 3A and 4D). In addition, simulations were repeated while including the nonspecific degradation rate, *μ*_ns_, and the increase in protein turnover resulted in a reduction in the recovery time and the duration of the phenotypic memory (Figure 4D). These results show that the delay between translational flux reallocation and reorganization of the proteome incurs a short term fitness cost by slowing adaptation, but confers a fitness advantage in fluctuating conditions as it allows cells to quickly return to optimal growth in the previous condition if the nutrient perturbation is short-lived. This phenotypic memory is also predicted to occur during starvation (Supplemental Figure 5), and is seen experimentally [40].

**Figure 4.**
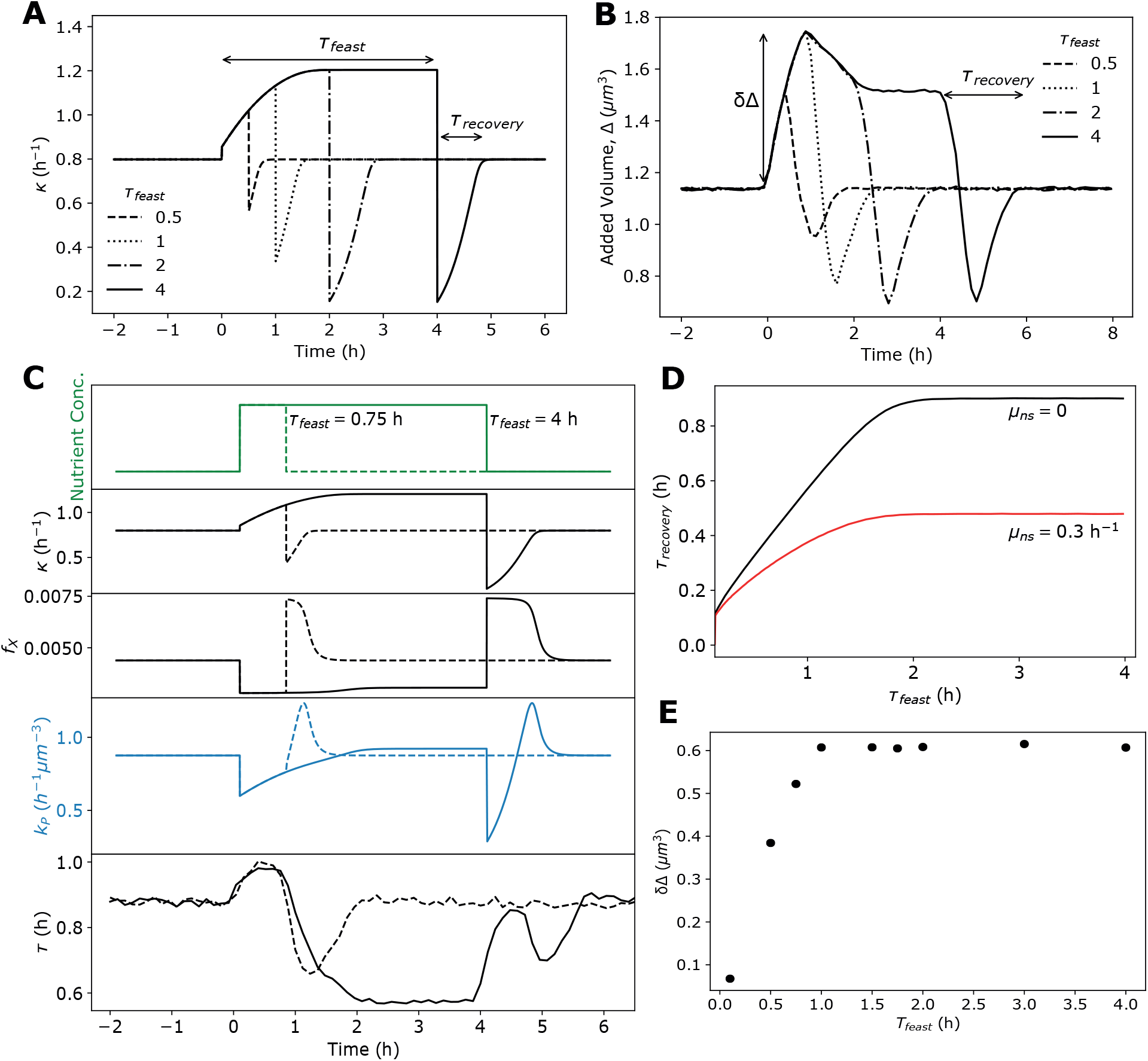
Proteome reallocation and cell size regulation in pulsatile nutrient environment. (A) Average singlecell growth rate simulations of bacteria experiencing a nutrient-rich pulse of duration *τ*_feast_. For each trajectory with pulse-length *τ*_feast_, the time required following downshift for the growth rate to return to within 99% of the pre-shift level was measured, given by *τ*_recovery_. (B) Average dynamics of added volume, Δ, for 300 single-cell trajectories experiencing a nutrient-rich pulse as shown in (A). The time required to stabilize at the initial added volume after pulse cessation is again given by *τ*_recovery_. (C) Example dynamics of simulation where *τ*_feast_ = 0.75 h, and *τ*_feast_ = 4 h. In both cases, the top four panels are deterministic simulations of average intracellular dynamics, whereas the bottom panel is the average dynamics of 300 single-cell stochastic simulations. (D) Quantification of the relationship of *τ*_feast_ and *τ*_recovery_ from the simulations in (A) for two different degradation rates. (E) Quantification of the relationship of *τ*_feast_ and *τ*_recovery_ from the simulations in (B). See Table I for a list of model parameters.

As with *τ*_recovery_, the overshoot in added volume, *δ*Δ, is also dependent on nutrient pulse length, such that increasing *τ*_feast_ increases *δ*Δ before saturating at a constant value (Figure 4E). This again is a consequence of the slow dynamics of proteome reallocation and stems from the prioritization of ribosome production over production of division machinery in response to nutrient upshift. This dynamic allocation strategy results in delayed division events, and thus an overshoot in added volume.

### Cell division is prioritized over biomass accumulation during nutrient downshift

Following the cessation of the nutrient-rich pulse, our model makes the interesting prediction that division protein synthesis is prioritized over ribosome production and biomass accumulation during downshift, as allocation to division protein synthesis transiently becomes maximal at the expense of ribosome allocation (Figure 1D). This behavior can be understood by recalling that *f_X_* and *f_P_* are co-regulated, and that an increase in *f_X_* necessarily requires a decrease in *f_R_* (Eq. 3). As a result, there is temporary increase in division rate (undershoot in *τ*) caused by an overshoot in *k_p_* (Figure 4C), while biomass accumulation temporarily slows (undershoot in *κ* Figure 4C), leading to a rapid reduction in cell size (undershoot in Δ, Figure 4B). This prioritization of division protein synthesis is a surprising prediction given that following downshift cells are experiencing harsher environmental conditions. We propose explanations for this behavior in the Discussion section.

Remarkably, our model predicts distinctly different recovery behavior in interdivision time following cessation of the nutrient-rich pulse, which is dependent on the time period of the nutrient pulse. This can be seen clearly by comparing the simulation dynamics of bacteria experiencing nutrient-rich pulses of 0.75 and 4 hrs (bottom panel, Figure 4C). Specifically, cells experiencing longer pulse lengths exhibit a non-monotonic recovery of interdivision time, *τ*, which is not observed at shorter pulse durations. This behavior can be understood by considering the impacts of both the overall translational flux (*J_t_* = *κ*) and the division protein allocation fraction (*f_X_*) on division protein synthesis rate, given by *k_P_* = *γJ_t_f_X_*. Under both conditions, *f_X_* behaves similarly immediately following downshift, namely that allocation to division protein production transiently increases before relaxing to its steady-state value (third panel from top, Figure 4C). As *k_P_* is proportional to *f_X_*, at short pulse lengths the increase in *f_X_* causes an overshoot in *k_P_* following downshift (fourth panel from top, Figure 4C). Importantly, there is a temporary undershoot in growth rate following downshift under both conditions, however the magnitude of this growth rate undershoot is significantly larger at longer pulse lengths (second panel from top, Figure 4C) due to a greater mismatch in metabolic and translational fluxes. As *k_P_* is also proportional to *κ*, at longer pulse lengths the initial drop in *k_P_* is due to a temporary halt of translation. This is followed by a translation flux ramp-up in which division is prioritized, resulting in a temporary overshoot in *k_P_*, and an overall non-monotonic recovery behavior in *τ*. Importantly, when the quality of the nutrient-rich media is reduced but the pulse length remains long, there is a reduced growth rate undershoot following pulse cessation, and the non-monotonic recovery in *τ* is not seen (Supplemental Figure 6). Thus, this pulse length-dependent division control is a direct consequence of the dependence of *k_P_* on both *f_X_* and *κ*.

### Cell size-dependent protein synthesis regulates recovery from stationary phase under pulsed nutrient supply

When the environmental nutrient supply has been exhausted, bacteria halt growth and enter stationary phase. Bacterial division control and size homeostasis behavior is markedly different in stationary phase compared to exponential phase, and a robust mechanistic model which captures size control dynamics in both phases of growth is still lacking. As such, we were interested if our model for dynamic proteome allocation would successfully predict cell size and division control upon exit from stationary phase. Under such conditions, the effects of protein turnover on cell physiology become crucial [41]. From Eq. (1) we see that although bacterial growth vanishes in stationary phase (*κ* = 0), protein production does not cease completely, but is balanced by the degradation rate such that the translational flux is given by 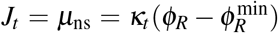. This implies that a small fraction of ribosomes remain active and that amino acid supply comes solely from protein turnover. Importantly, division protein production scales with cell volume and persists in stationary phase, with *k_P_* = *γf_X_μ*_ns_. As a result, the concentration of division proteins, *c_X_*, at steady-state in stationary phase is set by the relative rates of protein production and degradation, namely *c_X_* = *k_P_X*_0_/*μ_X_*, predicting that cells maintain a constant concentration of division proteins in stationary phase, regardless of cell size. Because division is dependent on the total number of division proteins and not its concentration, we therefore expect larger cells to divide faster upon nutrient exposure.

To examine if our model was able to capture cell division dynamics in different growth regimes, we simulated cell size dynamics of bacterial cells exiting stationary phase (i.e. starting at steady-state when *κ* = 0 and *c* = 0) through pulsatile nutrient exposure of constant duration with a variable separation time, *τ*_pulse_ (Figure 5A). As an increase in available nutrients results in an increase in the intracellular amino acid mass fraction (Supplemental Figure 7) [24], our model predicts that bacteria transiently prioritize ribosome production over division immediately following pulse exposure, similar to nutrient upshift behavior predicted in exponential phase (Figure 5A, Figure 3, Supplemental Figure 7). Consequently, immediately following nutrient influx, *k_P_* drops and the degradation rate dominates, resulting in a sharp decrease in the division protein number, *X*. Importantly, in the time between pulses, *X* increases significantly due to an increase in the division protein production rate caused by an increase in cell volume. This stands in contrast to a previous model for division control in stationary phase [24], which assumed that bacteria immediately allocate resources to division during nutrient upshift, causing the division protein production rate to transiently increase before falling to some basal value if the pulse rate is of insufficient frequency. Despite the stark differences in molecular details between these models, we find that the time from pulse onset to first division, *T*_lag_ (Figure 5B), monotonically decreases with increasing feedrate (decreasing *τ*_pulse_, Figure 5C), which is observed experimentally [24]. This behavior occurs because although bacteria initially prioritize ribosome production over division when exiting stationary phase, once the ribosome bottleneck is relieved, cells then upregulate division machinery. As a faster feedrate relieves this bottleneck quicker, a faster feedrate results in a shorter lag time until division. Also consistent with experimental results [24], we find that increasing the division protein degradation rate (*μ_x_*) increases *T*_lag_, while increasing the protein production rate (*k_P_*) decreases *T*_lag_ (Figure 5C), highlighting the importance of the degradation and volume-specific protein synthesis rates in controlling division timing.

**Figure 5.**
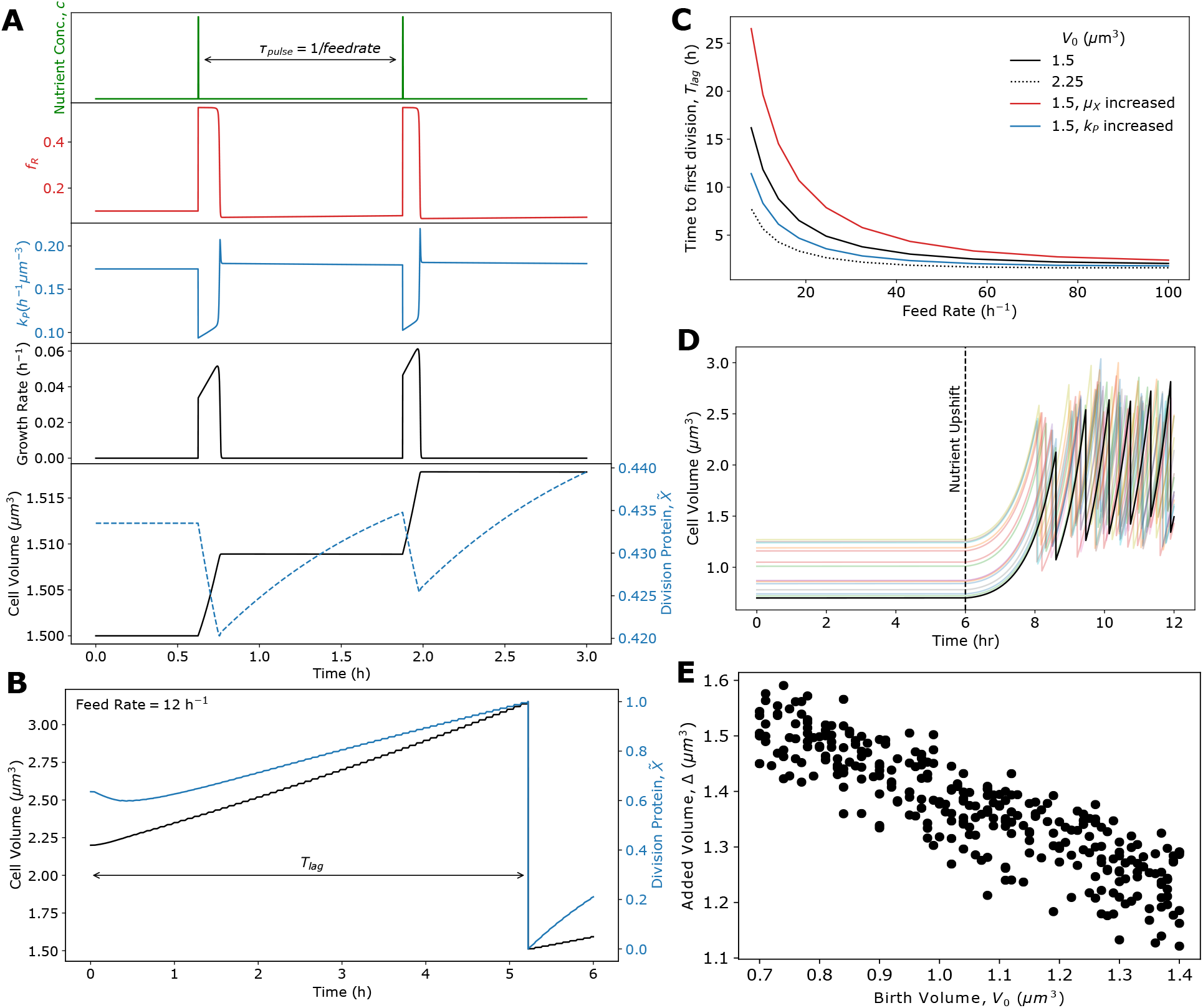
Cell size and division control during exit from stationary phase. (A) Single-cell simulation dynamics of ribosome allocation fraction, division protein production, growth rate, volume, and normalized X protein abundance for *E. coli* experiencing pulses of nutrients with delay *τ*_pulse_ starting from stationary phase. In response to an influx of nutrients, the cell temporarily decreases kp in order to produce ribosomes. (B) Cell volume and normalized division protein abundance dynamics for a feed rate of 12 h^-1^. (C) Using the simulation setup shown in (A), the time from pulsing onset until the first division event, *T*_lag_ (example trajectory shown in (B)), was measured as a function of pulse frequency (feed rate) for several initial volumes, degradation rates, and division protein production rates. For increased degradation, *μ_X_* = 1 h^-1^. For increased *k_P_, γα* = 2.875 *μm*^-3^ and *γβ* = 0.875 *μm*^-3^. (D) Example singlecell trajectories of cells with randomized initial volumes exited stationary phase via a single nutrient shift (dotted line). (E) Negative correlation between birth volume and added volume shows that *E. coli* exhibit sizer dynamics when exiting stationary phase, which is in agreement with experimental observations [42]. See Table I for a list of model parameters.

As cells in stationary phase maintain a constant concentration of division proteins regardless of size, our model predicts that *T*_lag_ is dependent on initial volume in stationary phase, *V*_0_, such that larger cells divide faster (Figure 5C). Importantly, this dependence of division timing on initial cell size is seen experimentally [42], and is not captured by the model proposed in Ref. [24]. To more specifically investigate size control mechanisms when exiting stationary phase, we simulated single-cell volume trajectories of bacteria exiting stationary phase via a single nutrient upshift (Figure 5D; see Supporting Information Section III). Importantly, we found that the adder model for cell size control did not hold under this growth regime, but rather cells exhibited sizer-like behavior, which is characterized by the added volume being negatively correlated with birth volume (Figure 5E). This behavior has been observed experimentally [42], and again can be understood from our threshold accumulation model, now considering the limit when *μ*_X_ ≫ *κ*. In such environments, bacteria divide once reaching a set size given by *V_d_* = *μ_X_/k_P_*.

## Discussion

We have developed a coarse-grained proteome sector model which quantitatively captures experimentally observed growth rate and size control dynamics in response to nutrient upshift in both exponential (Figure 3) and stationary phases (Figure 5). Our model highlights an important resource allocation trade-off that cells must make between optimizing for biomass accumulation or division in dynamic nutrient environments. In response to nutrient upshift, we predict that bacteria prioritize ribosome production in both exponential and stationary phase, resulting in faster biomass accumulation but delayed division. At the single-cell level, this results in a transient overshoot in both added volume and interdivision time. Interestingly, when simulating population level growth dynamics (see Supporting Information Section IV), we find that upshift results in a temporary reduction in population growth rate (Figure 6). This raises the question, in response to increased nutrient availability, why do bacteria temporarily slow proliferation? One possible explanation is that by delaying division, cellular resources are freed up which can be reallocated to quickly alleviate the growth bottleneck caused by a lack of ribosomes. As a result, cells are optimized for biomass accumulation instead of population growth, which allows for individual cells to adapt quickly to new environments. A second explanation is that because bacteria can quickly inactivate ribosomes [22] and recycle the amino acids through degradation, cells prioritize ribosome production as a method of energy storage when the environment is transiently nutrient-rich. Thus by producing ribosomes in response to nutrient upshift, bacteria simultaneously relieve the growth bottleneck caused by lack of ribosomes, while also being able to quickly convert metabolites into proteins which can be reallocated in the future after the nutrients have been exhausted. This strategy could allow for bacterial survival in harsher fluctuating environments, when nutrients are few and far between.

**Figure 6.**
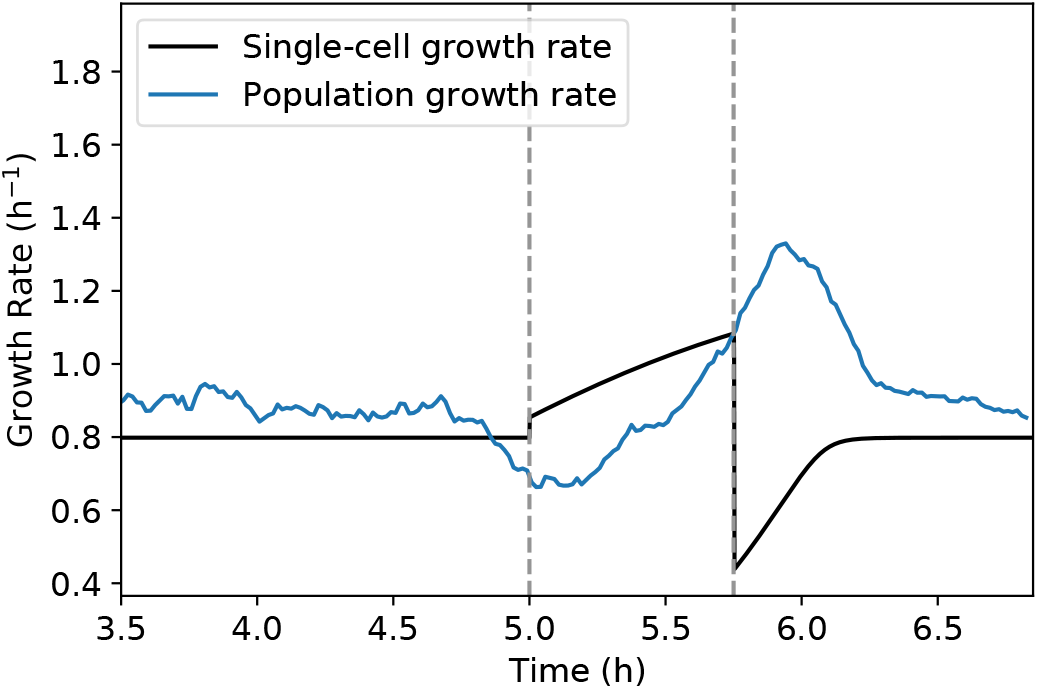
Cell proliferation dynamics during nutrient up-shift. Comparison of single-cell growth rate (black) and population growth rate (blue) dynamics in response to nutrient-rich pulse, where the population growth rate is given by the number of cell divisions per time. Dotted lines correspond to the start and end of the nutrient-rich period. See Table I for a list of model parameters.

With our model able to capture nutrient upshift dynamics, we simulated bacterial growth rate and size control dynamics in response to pulsatile nutrient exposure to predict how resources are allocated in more complicated time-varying environments. In such conditions, growth rate recovery time following nutrient downshift, increased with increasing pulse length (Figure 4), showing that bacteria exhibit a transient memory of the previous metabolic state. This phenotypic memory arises from the slow dynamics of proteome reallocation, and although it incurs a short term fitness cost, this passive mechanism can confer a fitness advantage in fluctuating conditions, as it allows cells to quickly return to optimal growth in the previous condition if the nutrient perturbation is short-lived.

Our simulations also yielded surprising predictions for the size control dynamics following cessation of nutrient-rich exposure (Figure 4C). In particular, our model predicts that bacteria transiently prioritize production of division proteins over production of ribosomes, resulting in a temporary undershoot in interdivision time and added volume. This result in striking, because it predicts that in response to onset of harsher environmental conditions, bacteria transiently upregulate the production of costly division machinery instead of prioritizing energy storage. In addition, this prioritization of division results in a temporary overshoot in population growth rate (Figure 6), meaning that the number of cells that must compete with each other for nutrients sharply increases in the new, less-favorable, environment. Several potential explanations for this behavior warrant exploration in future experimental and theoretical studies. First, by increasing division events, cells rapidly decrease cell size and thus increase surface-to-volume ratio [3, 43]. As a higher surface-to-volume ratio results in greater nutrient influx [44, 45], decreasing cell size may confer an important fitness advantage despite the metabolic coast associated with upregulating division protein production. Second, bacteria may employ this increased rate of division as a population bet-hedging strategy which facilitates adaptation to fluctuating environments. Previous work has shown that partitioning of cellular contents at division is a major determinant of phenotypic heterogeneity [46]. Thus, by transiently increasing the number of division events, a bacterial population temporarily will exhibit a broader range of phenotypes. Phenotypic heterogeneity increases in adverse environments in both prokaryotic and eukaryotic populations, and previous work has shown that heterogeneity promotes adaptation to time-varying stress by facilitating development of resistance-conferring mutations and/or by alleviating the fitness cost of constitutive expression of unnecessary proteins [47–50]. These results suggest that bacterial cells utilize division control to increase population heterogeneity in response to harsh environmental perturbations, thus facilitating adaptation to new environments and conferring increased population fitness in time-varying environments.

## Author Contributions

J.C.K. and S.B. designed and developed the study. J.C.K. carried out the simulations and analyzed the data. J.C.K. and S.B. wrote the article.

## Acknowledgements

This work was supported by National Institute of General Medical Sciences of the National Institutes of Health under award numbers R35 GM143042 (SB), 5T32 GM133353 (JCK), and the Shurl and Kay Curci Foundation (SB).

## Notes

### Competing Interest Statement

The authors have declared no competing interest.

### Summary of Updates

Revised authorship order.

## References

[1] S. Jun, F. Si, R. Pugatch, and M. Scott, Reports on Progress in Physics 81, 056601 (2018).

[2] D. Serbanescu, N. Ojkic, and S. Banerjee, FEBS Journal (2021), 10.1111/febs.16234.

[3] L. K. Harris and J. A. Theriot, Cell 165, 1479 (2016).

[4] M. Mori, S. Schink, D. W. Erickson, U. Gerland, and T. Hwa, Nature Communications 8, 1 (2017).

[5] M. Panlilio, J. Grilli, G. Tallarico, I. Iuliani, B. Sclavi, P. Cicuta, and M. C. Lagomarsino, Proceedings of the National Academy of Sciences 118 (2021), 10.1073/PNAS.2016391118.

[6] J. Nguyen, V. Fernandez, S. Pontrelli, U. Sauer, M. Ackermann, and R. Stocker, Nature Communications 12, 1 (2021).

[7] M. Schaechter, O. Maaløe, and N. Kjeldgaard, Microbiology 19, 592 (1958).

[8] F. Si, D. Li, S. E. Cox, J. T. Sauls, O. Azizi, C. Sou, A. B. Schwartz, M. J. Erickstad, Y. Jun, X. Li, and S. Jun, Current Biology 27, 1278 (2017).

[9] H. Zheng, Y. Bai, M. Jiang, T. A. Tokuyasu, X. Huang, F. Zhong, Y. Wu, X. Fu, N. Kleckner, T. Hwa, and C. Liu, Nature Microbiology 5, 995 (2020).

[10] D. Serbanescu, N. Ojkic, and S. Banerjee, Cell Reports 32, 108183 (2020).

[11] M. Campos, I. V. Surovtsev, S. Kato, A. Paintdakhi, B. Beltran, S. E. Ebmeier, and C. Jacobs-Wagner, Cell 159, 1433 (2014).

[12] S. Taheri-Araghi, S. Bradde, J. T. Sauls, N. S. Hill, P. A. Levin, J. Paulsson, M. Vergassola, and S. Jun, Current Biology 25, 385 (2015).

[13] S. Hui, J. M. Silverman, S. S. Chen, D. W. Erickson, M. Basan, J. Wang, T. Hwa, and J. R. Williamson, Molecular systems biology 11, 784 (2015).

[14] A. Schmidt, K. Kochanowski, S. Vedelaar, E. Ahrné, B. Volkmer, L. Callipo, K. Knoops, M. Bauer, R. Aebersold, and M. Heinemann, Nature Biotechnology 34, 104 (2016).

[15] M. Mori, Z. Zhang, A. Banaei-Esfahani, J. Lalanne, H. Okano, B. C. Collins, A. Schmidt, O. T. Schubert, D. Lee, G. Li, R. Aebersold, T. Hwa, and C. Ludwig, Molecular Systems Biology 17 (2021), 10.15252/msb.20209536.

[16] M. Scott, C. W. Gunderson, E. M. Mateescu, Z. Zhang, and T. Hwa, Science 330, 1099 (2010).

[17] M. Basan, M. Zhu, X. Dai, M. Warren, D. Sévin, Y. Wang, and T. Hwa, Molecular Systems Biology 11, 836 (2015).

[18] D. W. Erickson, S. J. Schink, V. Patsalo, J. R. Williamson, U. Gerland, and T. Hwa, Nature 551, 119 (2017).

[19] F. Bertaux, J. Von Kügelgen, S. Marguerat, and V. Shahrezaei, PLoS Computational Biology 16, e1008245 (2020).

[20] N. M. Belliveau, G. Chure, C. L. Hueschen, H. G. Garcia, J. Kondev, D. S. Fisher, J. A. Theriot, and R. Phillips, Cell Systems 12, 924 (2021).

[21] B. D. Towbin, Y. Korem, A. Bren, S. Doron, R. Sorek, and U. Alon, Nature Communications 8, 1 (2017).

[22] X. Dai, M. Zhu, M. Warren, R. Balakrishnan, V. Patsalo, H. Okano, J. R. Williamson, K. Fredrick, Y. P. Wang, and T. Hwa, Nature Microbiology 2 (2016), 10.1038/nmicrobiol.2016.231.

[23] G. Lambert and E. Kussel, PLoS Genetics 10 (2014), 10.1371/journal.pgen.1004556.

[24] K. Sekar, R. Rusconi, J. T. Sauls, T. Fuhrer, E. Noor, J. Nguyen, V. I. Fernandez, M. F. Buffing, M. Berney, S. Jun, R. Stocker, and U. Sauer, Molecular Systems Biology 14, e8623 (2018).

[25] K. R. Ghusinga, C. A. Vargas-Garcia, and A. Singh, Scientific Reports 6, 1 (2016).

[26] F. Si, G. Le Treut, J. T. Sauls, S. Vadia, P. A. Levin, and S. Jun, Current Biology 29, 1760 (2019).

[27] M. Scott, S. Klumpp, E. M. Mateescu, and T. Hwa, Molecular Systems Biology 10, 747 (2014).

[28] D. Molenaar, R. Van Berlo, D. De Ridder, and B. Teusink, Molecular Systems Biology 5, 323 (2009).

[29] P. P. Pandey and S. Jain, Theory in Biosciences 135, 121 (2016).

[30] N. Giordano, F. Mairet, J. L. Gouzé, J. Geiselmann, and H. de Jong, PLoS Computational Biology 12, 1 (2016).

[31] Y. K. Kohanim, D. Levi, G. Jona, B. D. Towbin, A. Bren, U. A. Correspondence, and U. Alon, Cell Reports 23 (2018), 10.1016/j.celrep.2018.05.007.

[32] J. S. Edwards, R. U. Ibarra, and B. O. Palsson, Nature Biotechnology 19, 125 (2001).

[33] R. U. Ibarra, J. S. Edwards, and B. O. Palsson, Nature 420, 186 (2002).

[34] N. E. Lewis, K. K. Hixson, T. M. Conrad, J. A. Lerman, P. Charusanti, A. D. Polpitiya, J. N. Adkins, G. Schramm, S. O. Purvine, D. Lopez-Ferrer, et al., Molecular Systems Biology 6, 390 (2010).

[35] L. U. Magnusson, A. Farewell, and T. Nyström, Trends in Microbiology 13, 236 (2005).

[36] C. Wu, R. Balakrishnan, N. Braniff, M. Mori, G. Manzanarez, Z. Zhang, and T. Hwa, Proceedings of the National Academy of Sciences of the United States of America 119, 1 (2022).

[37] A. M. Fallon, C. S. Jinks, G. D. Strycharz, and M. Nomura, Proceedings of the National Academy of Sciences of the United States of America 76, 3411 (1979).

[38] L. Herzel, J. A. Stanley, C.-C. Yao, and G.-W. Li, Nucleic Acids Research 50, 5029 (2022).

[39] M. Zhu, M. Mori, T. Hwa, and X. Dai, Nature Microbiology 4, 2347 (2019).

[40] A. Jõers and T. Tenson, Scientific Reports 6, 1 (2016).

[41] L. Calabrese, J. Grilli, M. Osella, C. P. Kempes, M. C. Lagomarsino, and L. Ciandrini, PLoS Computational Biology 18, e1010059 (2022).

[42] S. Bakshi, E. Leoncini, C. Baker, S. J. Cañas-Duarte, B. Okumus, and J. Paulsson, Nature Microbiology 6, 783 (2021).

[43] N. Ojkic, D. Serbanescu, and S. Banerjee, eLife 8, e47033 (2019).

[44] K. D. Young, Microbiology and Molecular Biology Reviews 70, 660 (2006).

[45] N. Ojkic and S. Banerjee, Biophysical Journal 120, 2079 (2021).

[46] P. Thomas, G. Terradot, V. Danos, and A. Y. Weiße, Nature Communications (2018), 10.1038/s41467-018-06912-9.

[47] E. Kussell and S. Leibler, Science 309, 2075 (2005).

[48] D. Van Dijk, R. Dhar, A. M. Missarova, L. Espinar, W. R. Blevins, B. Lehner, and L. B. Carey, Nature Communications 6 (2015), 10.1038/ncomms8972.

[49] B. M. Martins and J. C. Locke, Current Opinion in Microbiology 24, 104 (2015).

[50] Z. Bódi, Z. Fárkás, D. Nevozhay, D. Kalapis, V. Lazar, B. Csörgő, A. Nyerges, B. Szamecz, G. Fekete, B. Papp, H. Araújo, J. L. Oliveira, G. Moura, M. A. Santos, T. Székely, G. Balázsi, and C. Pál, PLoS Biology 15, 1 (2017).

