## Supporting Information for "Dynamic trade-offs between biomass accumulation and division determine bacterial cell size and proteome in fluctuating nutrient environments"

### I. COARSE-GRAINED MODEL FOR BACTERIAL GROWTH AND SIZE CONTROL

**Model derivation.** Here we present a four-component coarse-grained proteome sector model to describe cellular growth rate and division control. These sectors consist of housekeeping proteins (Q), ribosomal proteins (R), metabolic proteins (P), and division proteins (X). Bacterial cells grow exponentially in size during the cell cycle, such that the dynamics of cell volume, $V$, are given by

$$\frac{\mathrm{d}V}{\mathrm{d}t} = \kappa V \;, \tag{1}$$

where $\kappa$ is the growth rate of the cell. Assuming constant protein density [1], growth rate can also be expressed in terms of total protein mass, $M$, such that

$$\kappa = \frac{1}{M}\frac{dM}{dt} \;. \tag{2}$$

The rate of change of protein mass is proportional to the mass of actively translating ribosomes. In addition, if we assume that protein turnover is governed by a constant, nonspecific degradation rate $\mu_{\mathrm{ns}}$, then the rate rate of change of protein mass is given by

$$\frac{\mathrm{d}M}{\mathrm{d}t} = \kappa_t(M_R - M_R^{\mathrm{in}}) - \mu_{\mathrm{ns}}M \;, \tag{3}$$

where $\kappa_t$ is the translational efficiency of the cell, $M_R$ is the total mass of ribosomes, and $M_R^{\mathrm{in}}$ is the mass of inactive ribosomes. Using Eq. (2), the growth rate can then be defined as

$$\kappa = \kappa_t(\phi_R - \phi_R^{\mathrm{min}}) - \mu_{\mathrm{ns}} \;. \tag{4}$$

where $\phi_R = M_R/M$ is the ribosome mass fraction and $\phi_R^{\mathrm{min}} = M_R^{\mathrm{in}}/M$ is the mass fraction of inactive ribosomes. To derive the dynamics of $\phi_R$, we note that

$$\frac{\mathrm{d}M_R}{\mathrm{d}t} = \kappa_t f_R(M_R - M_R^{\mathrm{in}}) - \mu_{\mathrm{ns}}M_R \;, \tag{5}$$

where $f_R$ is the fraction of total cellular protein synthesis flux devoted to ribosomes. It follows that the time dynamics of $\phi_R$ are then

$$\frac{\mathrm{d}\phi_R}{\mathrm{d}t} = \kappa_t(a)(\phi_R - \phi_R^{\mathrm{min}})(f_R(a) - \phi_R) \;. \tag{6}$$

Both $\kappa_t$ and $f_R$ depend on the amino acid concentration in the cell, which in turn depends on the nutrient availability. We note that the inclusion of a nonspecific degradation term does not impact the dynamics of $\phi_R$, as all cellular proteins are assumed to be degraded at the same rate. To connect cellular growth rate to amino acid mass fraction ($a$) and nutrient availability we use the following condition for flux balance [2]:

$$\frac{\mathrm{d}a}{\mathrm{d}t} = \kappa_n(a)\phi_P - \kappa \;, \tag{7}$$

where $\kappa_n$ is the nutritional efficiency of the cell and $\phi_P$ is the mass fraction of P-sector protein that are responsible for transporting nutrients into the cell. As $\phi_Q$ is invariant to nutrient perturbations, the mass fractions of each sector satisfy the constraint $\phi_R + \phi_P + \phi_X = 1 - \phi_Q = \phi^{\max}$. Using this relation, along with our definition of growth rate from Eq. (4), the amino acid mass fraction can be rewritten as

$$\frac{\mathrm{d}a}{\mathrm{d}t} = \kappa_n(a)(\phi^{\max} - \phi_R - \phi_X) - \kappa_t(a)(\phi_R - \phi_R^{\min}) + \mu_{\mathrm{ns}} \; . \tag{8}$$

We note that amino acid supply is now given by a combination of nutrient import and amino acid recycling due to protein turnover. To make explicit the dependency of the efficiencies, $\kappa_n$ and $\kappa_t$, on $a$, we define two regulatory functions, $f(a)$ and $g(a)$, as given by [3]. Specifically, we assume $\kappa_n = \kappa_n^0(c) f(a)$ and $\kappa_t = \kappa_t^0 g(a)$, where $\kappa_t^0$ is a constant, and $\kappa_n^0$ is a function of the extracellular nutrient concentration $c$. The regulatory functions are then:

$$f(a) = \frac{1}{1 + (a/a_n)^2} \; , \tag{9}$$

$$g(a) = \frac{(a/a_t)^2}{1 + (a/a_t)^2} \; , \tag{10}$$

where translation becomes significantly attenuated for amino acid concentrations below $a_t$, and the amino acid supply flux becomes significantly attenuated by feedback inhibition for $a$ above $a_n$.

Similar to the dynamics of $\phi_R$, to derive the dynamics of $\phi_X$ we begin with the equation for the mass of protein X

$$\frac{\mathrm{d}M_X}{\mathrm{d}t} = \kappa_t f_X (M_R - M_R^{\mathrm{in}}) - \mu_X M_X \; , \tag{11}$$

where $f_X$ is the fraction of total synthesis capacity of a cell devoted to making cell division proteins, and $\mu_X$ is the degradation rate of protein X. This degradation rate is specific to protein X and can be different than the nonspecific degradation rate $\mu_{\mathrm{ns}}$. This leads to the following equation for $\phi_X$

$$\frac{\mathrm{d}\phi_X}{\mathrm{d}t} = \kappa_t(\phi_R - \phi_R^{\min})(f_X - \phi_X) - (\mu_X - \mu_{\mathrm{ns}})\phi_X \; . \tag{12}$$

When $\mu_X \approx \mu_{\mathrm{ns}}$, the last term on the right hand side of Eq. (12) is negligible and the equation takes the same form as Eq. (6), allowing the dynamics of each sector to be written in vector form, as presented in the main text in Eq. (2). To derive the time evolution of the amount of protein X, Eq. (12) can be rewritten in terms of the concentration of $X$, $c_X = \phi_X \rho_c / m_X$, where $\rho_c$ is the protein mass density of the cell and $m_X$ is the mass of molecule X. We then get

$$\frac{\mathrm{d}c_X}{\mathrm{d}t} = \frac{f_X \rho_c}{m_X}(\kappa + \mu_{\mathrm{ns}}) - c_X(\kappa + \mu_X) \; , \tag{13}$$

remembering that $\kappa = \kappa_t(\phi_R - \phi_R^{\min}) - \mu_{\rm ns}$. If the total amount of division proteins is $X = c_X V$, then using Eqs. (1) and (13) we can obtain the time dynamics of $X$, where

$$\frac{\mathrm{d}X}{\mathrm{d}t} = c_X \frac{\mathrm{d}V}{\mathrm{d}t} + \frac{\mathrm{d}c_X}{\mathrm{d}t} V \tag{14}$$

$$= \frac{f_X \rho_c}{m_X}(\kappa + \mu_{\rm ns})V - \mu_X X \; . \tag{15}$$

In this model, division is triggered at $t = \tau$ after the cell has accumulated a fixed number of X proteins such that $X(\tau) = X_0$. For simplicity, we normalize Eq. (15) by $X_0$ to yield the time dynamics of the fraction of the total number of division proteins required to trigger cell division, such that

$$\frac{\mathrm{d}X/X_0}{\mathrm{d}t} = \frac{\mathrm{d}\tilde{X}}{\mathrm{d}t} = \gamma f_X(\kappa + \mu_{\rm ns})V - \mu_X \tilde{X} \; , \tag{16}$$

where $\gamma = \rho_c / X_0 m_X$ and here now cells divide when $\tilde{X}(\tau) = 1$. Eq. (16) demonstrates that division proteins are synthesized proportionally to the cell volume, such that the concentration remains constant at at given growth rate regardless of cell volume. This allows us to identify the division protein synthesis rate per unit volume, $k_P$, given by

$$k_P = \gamma f_X(\kappa + \mu_{\rm ns}) \; . \tag{17}$$

As with the mass fractions, the allocation fractions are constrained such that $f_R + f_P + f_X = 1 - f_Q = \phi^{\max}$, meaning that two regulatory functions must be defined in order to simulate the dynamics of growth rate and cell size control. To do so, we assume that allocation is dependent on the amino acid pool, and that the division protein sector (X) is partially co-regulated with the metabolic protein sector (P), such that $f_X(a)$ is given by a linear combination of two sub-sectors: $f_X^\alpha(a)$, which denotes the portion which is co-regulated with the metabolic sector, and $\beta$, which denotes the basal allocation fraction. If we define $f_P^*(a) = f_P(a) + f_X^\alpha(a)$ as the proteome fraction which is co-regulated opposite of $f_R(a)$, then $f_X(a)$ can be expressed as a function of $f_R(a)$, such that

$$f_X(a) = f_X^\alpha(a) + \beta = \alpha f_P^*(a) + \beta = \alpha(\phi_R^{\max} - f_R(a)) + \beta \; , \tag{18}$$

where $\alpha$ is the fraction of $f_P^*$ which contains the co-regulated portion of division proteins and where $\phi_R^{\max} = \phi^{\max} - \beta$. The fraction of total synthesis capacity devoted to production of ribosomes, $f_R(a)$, is given by

$$f_R(a) = \frac{-f'(a)g(a)\phi_R^{\max} + f(a)g'(a)\phi_R^{\min}}{-f'(a)g(a) + f(a)g'(a)} \; , \tag{19}$$

in which $f_R$ is chosen to maximize amino acid flux at steady state (derived in Section II).

With $f_X$ now defined, the division protein synthesis rate can now be rewritten in terms of $f_R$, where

$$k_P = \gamma(\alpha(\phi_R^{\max} - f_R(a)) + \beta)(\kappa + \mu_{\rm ns}) \; . \tag{20}$$

At steady state $k_P$ can be rewritten solely as a function of growth rate, such that

$$k_P(\kappa) = \gamma(\alpha(\Delta\phi - \frac{\kappa + \mu_{\rm ns}}{\kappa_t}) + \beta)(\kappa + \mu_{\rm ns}) , \tag{21}$$

where $\Delta\phi = \phi_R^{\max} - \phi_R^{\min}$.

Two important features regarding division protein production can be seen from Eq. (21). First, division protein production as a function of growth rate is non-monotonic. At low growth rates, $k_P$ increases as $\kappa$ increases due to an increase in translational efficiency which results from an increase in amino acid availability. At higher growth rates, $k_P$ decreases as $\kappa$ increases due to a decrease in allocation of translational flux to division protein production (decrease in $f_X$). Second, due to the inclusion of a degradation rate, in stationary phase (when $\kappa = 0$), there remains a basal synthesis rate of division proteins which ensures that the division protein amount does not go to zero.

**Size control mechanism is dependent on growth conditions.** The coupled equations (1), (6), (8), and (16) now define the dynamics of the system. Assuming that at cell birth, the amount of division molecules is reset to zero (i.e. $\tilde{X}(0) = 0$), at steady state growth equations (1) and (16) can be solved to obtain

$$\tilde{X}(t) = \frac{k_P V_0}{\mu_X + \kappa}(e^{\kappa t} - e^{-\mu_X t}) , \tag{22}$$

where $V_0$ is cell volume at birth. If cells divide symmetrically (i.e. $V_d = 2V_0$) at $t = \tau$ after accumulating the required number of X proteins such that $X(\tau) = X_0$ or equivalently $\tilde{X}(\tau) = 1$, cell size at division can be related to $\kappa$ and $k_P$, yielding

$$1 = \frac{k_P}{\mu_X + \kappa}(V_d - V_0 2^{-\mu_X/\kappa}) , \tag{23}$$

where the cell volume at division is $V_d = V_0 e^{\kappa\tau}$. In the limit $\kappa >> \mu_X$, we arrive at

$$\Delta = \frac{\kappa}{k_P} , \tag{24}$$

where $\Delta = V_d - V_0$ is the added volume per generation. Because $k_P$ and $\kappa$ are both constant for a given growth medium, cells exhibit an adder mechanism in which a constant volume is added each generation regardless of birth size. In the opposite limit, in which $\kappa << \mu_X$, we get

$$V_d = \frac{\mu_X}{k_P} . \tag{25}$$

In slow-growing media, cells divide at a constant volume, and thus break from the adder mechanism and exhibit sizer behavior, in which cells divide at a set volume, regardless of birth volume.

**Connection to size law.** Equation (23) can be rearranged to give the birth size as a function growth rate, such that

$$V_0 = \frac{\kappa + \mu_X}{k_P(2 - 2^{-\mu_X/\kappa})} \,. \tag{26}$$

As $k_P$ is also a function of growth rate, Eq. (21) can be used to modify the equation above so that the dependency of $V_0$ on $\kappa$ is fully realized. As a result we obtain

$$V_0 = \frac{\kappa + \mu_X}{\gamma(\alpha(\Delta\phi - \frac{\kappa+\mu_{\text{ns}}}{\kappa_t}) + \beta)(\kappa + \mu_{\text{ns}})(2 - 2^{-\mu_X/\kappa})} \,. \tag{27}$$

Outside of slow growing conditions, the effects of protein degradation are negligible. Assuming $\kappa \gg \mu_{\text{ns}}$ and $\kappa \gg \mu_X$, the above equation simplifies to

$$V_0 = \frac{1}{\gamma\alpha(\Delta\phi - \kappa/\kappa_t) + \gamma\beta} \,. \tag{28}$$

This expression asymptotically approaches a maximum growth rate given by $\kappa_{\max} = \kappa_t(\Delta\phi + \beta/\alpha)$. It is important to note that this theoretical maximum is nonphysical as it assumes that $f_X = \phi_X = 0$, which is never the case given our definition of $f_X$. The actually maximum growth rate occurs when $\phi_R = \phi_R^{\max}$, thus giving an upper limit to physical growth rate at $\kappa_{\max} = \kappa_t \Delta\phi$. Equation (28) also predicts that there is no maximum cell size. However, our expression for $f_X$ constrains cell size to a finite value. When allocation to ribosomes is maximal, $\phi_X = \beta$, such that the maximum birth volume $V_0$ is given by $V_0^{\max} = 1/\gamma\beta$.

### II. GROWTH RATE MAXIMIZATION TO OBTAIN RIBOSOMAL ALLOCATION FRACTION

During steady-state exponential growth, the rate of amino acid supply is balanced by the rate of amino acid consumption through protein synthesis to ensure that there is no net change in the amino acid concentration. Furthermore, the rate of protein synthesis equals the rate of bacterial growth, so the cell is faced with the dual objectives of balancing and maximizing the amino acid flux in order to maximize the growth rate (Figure 1A,B). For a given translational efficiency $\kappa_t(a)$ and the nutritional efficiency $\kappa_n(a)$ (as determined by the growth medium), the organism must choose the ribosomal protein fraction $f_R(a)$ that balances the amino acid flux. This is implemented as follows [4]. At steady-state $\phi_R = f_R$ (Eq. 6). Using the condition of steady-state in Eq. (6) and (8) we get

$$f_R(a) = \frac{\kappa_n(a)\phi_R^{\max} + \kappa_t(a)\phi_R^{\min} + \mu_{\text{ns}}}{\kappa_n(a) + \kappa_t(a)} \,. \tag{29}$$

The steady-state growth rate is given by

$$\kappa(a) = \kappa_t(a)(f_R - \phi_R^{\min}) - \mu_{\text{ns}} \tag{30}$$

$$= \frac{\kappa_t(a)\Delta\phi - \mu_{\text{ns}}}{1 + \frac{\kappa_t(a)}{\kappa_n(a)}} \tag{31}$$

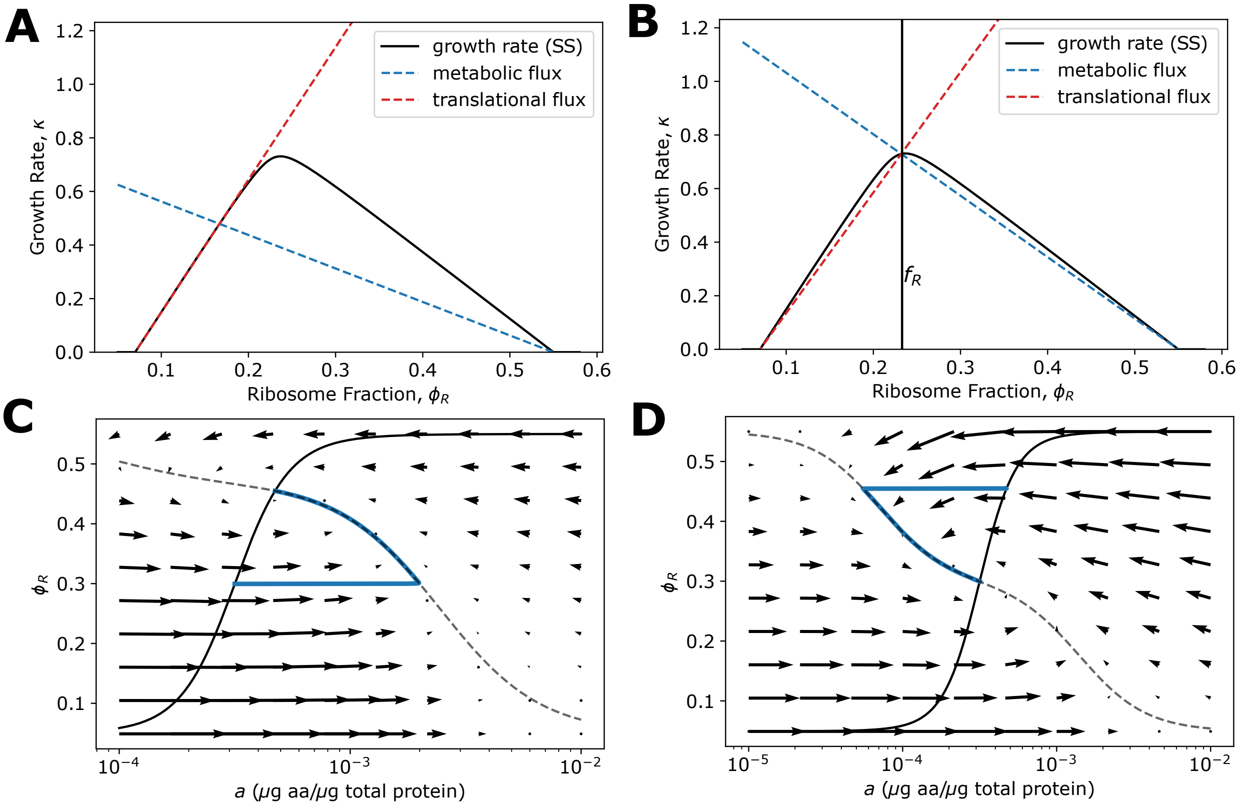


**Supplementary Figure** 1. **Flux balance by regulatory functions**. **A,B** Representative rate-balance plots of when translational and metabolic fluxes are balanced but growth is not maximal (**A**), and when fluxes are simulateneously balanced and maximized (**B**). $f_R$ is defined such that for a given nutrient condition, $f_R$ is equal to the ribosome fraction which both balances and maximizes flux. **C,D** Phase-plane analysis of model during nutrient upshift (**C**) and downshift (**D**). The model trajectory is shown in blue, showing behavior immediately after the nutrient shift occurs. The nullclines for $\phi_R$ and $a$ are shown as solid and dashed curves, respectively.

where $\Delta\phi = \phi_R^{\max} - \phi_R^{\min}$. Maximizing growth rate via $\kappa'(a) = 0$, we get the condition

$$\mu_{\rm ns} = \frac{\kappa_n^0 g'(a) f(a)^2 \Delta\phi + \kappa_t^0 g(a)^2 f'(a) \Delta\phi}{g(a) f'(a) - g'(a) f(a)} \,. \tag{32}$$

Substituting the above expression in the equation for $f_R$ we get the functional dependence of $f_R$ on amino acid concentration.

$$f_R(a) = \frac{\kappa_n^0 f(a) \phi_R^{\max} + \kappa_t^0 g(a) \phi_R^{\min} + \mu_{\rm ns}}{\kappa_n^0 f(a) + \kappa_t^0 g(a)} \tag{33}$$

$$= \frac{-f'(a) g(a) \phi_R^{\max} + f(a) g'(a) \phi_R^{\min}}{-f'(a) g(a) + f(a) g'(a)} \,. \tag{34}$$

**Note on dynamic flux balance and maximization.** Changes in available nutrient concentration, $c$, affect the metabolic flux through $\kappa_n$. As a result, in response to nutrient fluctuations, the cell must dynamically match the translational flux to the metabolic flux to prevent unsustainable build-up or depletion of the amino acid pool. The cell is able to alter both fluxes through three mechanisms which respond on two different timescales. The translational and nutritional efficiencies, $\kappa_t$ and $\kappa_n$, depend only on the current cellular amino acid levels (Eqs. (9) and (10)), and so can adjust quickly in response to changes in nutrient conditions. Following a nutrient shift, there is a sudden jump in $a$ caused by a temporary mismatch in fluxes. The regulatory functions ensure that the fluxes quickly rebalance, eliminating any subsequent dramatic changes in $a$ until the next change in $c$. The cell achieves further changes in flux by adjusting its ribosome mass fraction, $\phi_R$, via adjustment to the fraction of ribosomes allocated to synthesize additional ribosomes, $f_R$. Importantly, altering $\phi_R$ via $f_R$ is accomplished through protein synthesis and degradation, and thus occurs on a much slower timescale than changes to $\kappa_t$ and $\kappa_n$. This behavior can be seen graphically using phase-plane analysis, in which the model trajectory first moves to the $a$ nullcline by changing $a$ without changing $\phi_R$, at which point the fluxes have been rebalanced, before then moving to the fixed point by changing both $a$ and $\phi_R$ (Figure 1C,D).

### III. SIMULATING STOCHASTIC SINGLE CELL VOLUME TRAJECTORIES

In our modeling of growth rate, amino acid, and proteome allocation dynamics, we simulated deterministic trajectories by numerically solving the coupled ODEs defined by Eqs. (2) and (4) in the main text. These solutions predict the average single cell behavior. In order to investigate size control mechanisms, we must also include the growth rate and division noise observed in real biological systems. To this end, we also simulated single cell volume trajectories using a continuous-time stochastic hybrid system. In this setup, we introduce two sources of noise at the generational level. First, we consider growth rate noise. Although there is wide cell-to-cell variability in growth rate in the same nutrient environment, experimental data reveal that there is essentially no correlation between the growth rate of a mother and it's daughter cells [5]. Thus at steady-state, growth rate noise can be implemented by representing $\kappa$ as an independent random variable drawn from a normal distribution at cell birth [6]. We extend this framework into time-varying environments, and model growth rate noise in single cells by drawing an offset value, $\delta\kappa$, at the start of each new generation such that the growth rate for the $i$th cell is given by

$$\kappa_i(t) = \langle\kappa(t)\rangle + \delta\kappa_i\,, \tag{35}$$

where $\langle\kappa(t)\rangle$ is the population average growth rate at time $t$. Our analysis of single cell growth rate data [7] shows that the distribution of growth rates remains approximately Gaussian throughout nutrient upshift (Figure 2A), and that the standard deviation of the distribution, $\sigma_\kappa$ is a linear function of the growth rate, such that

$$\sigma_\kappa = a\langle\kappa\rangle + b \, , \tag{36}$$

where $a$ and $b$ are parameters obtained from fitting $\kappa$ vs $\sigma_\kappa$ data through the shift (Figure 2B,C). As a result, a single offset value, $\delta\kappa$, is drawn from a normal distribution with mean 0 and standard deviation $\sigma_\kappa$ at the start of each cell cycle, and remains constant until cell division. Using this cell-specific growth rate, each volume trajectory can be computed using

$$\frac{\mathrm{d}V_i}{\mathrm{d}t} = \kappa_i(t)V_i \, , \tag{37}$$

where this equation is coupled to Eq. (6) in the main text to determine cell division events. Cells do not divide exactly symmetrically, but instead exhibit partitioning error at division [8]. To implement this second source of noise into our simulation, the birth volume of the daughter cell, $V_0$, is given by the previous generation's final volume, $V_d$, multiplied by a random variable $r$, which is drawn from a normal distribution with mean 0.5 and standard deviation 0.04 [8], such that

$$V_0^{i+1} = rV_d^i \, , \tag{38}$$

where $i$ and $i+1$ denote the mother and daughter generations, respectively.

**Simulating stochastic trajectories when exiting stationary phase.** Bacterial cease biomass accumulation in stationary phase. As such, introducing growth rate noise at the generational level when simulating exit from stationary phase in the manner described above is not feasible. Instead, to probe cell division control in our model, we introduced noise in $\tilde{X}_0$, the threshold value required to trigger cell division. Specifically, for each cell a unique value of $\tilde{X}_0$ was drawn from a normal distribution with mean 1 and standard deviation 0.05.

### IV. SIMULATING STOCHASTIC POPULATION SIMULATIONS

Population level simulations were carried out using the same procedure as the single cell stochastic volume simulations, except for at each division event, both daughter cells were tracked, leading to a growing number of cell trajectories over time. Symmetric division noise was implemented similar to the single cell simulations, with one value of $r$ drawn at each division, yielding corresponding birth sizes for daughter cells 1 and 2 as

$$V_{0,1}^{i+1} = rV_d^i \, , \tag{39}$$

$$V_{0,2}^{i+1} = (1-r)V_d^i \, . \tag{40}$$

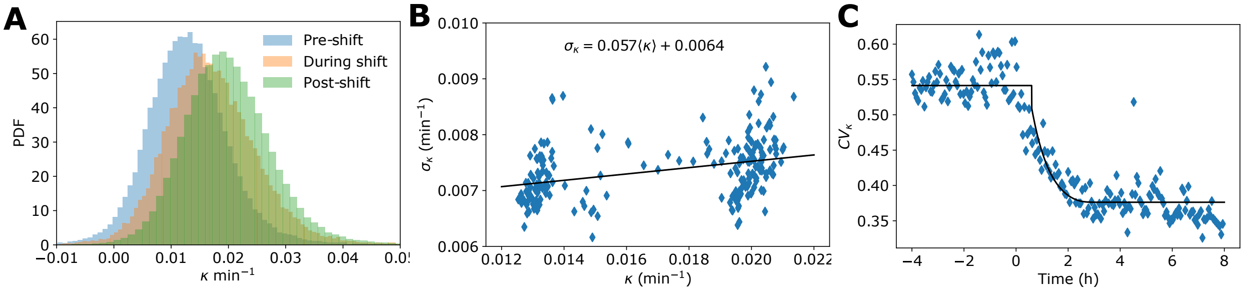


**Supplementary Figure** 2. **Simulating growth rate noise during nutrient upshift**. **A** Single cell growth rate distributions remains approximately Gaussian during nutrient upshift. Growth rate data from Ref. [7] are binned based on time of division into Pre-shift (-4-0 hrs), During shift (0-2.5 hrs), and Post-shift (2.5-8 hrs). **B** The standard deviation remained proportional to the average single cell growth rate during nutrient upshift, allowing time-binned growth rate ($\kappa$) vs standard deviation ($\sigma_\kappa$) data to be fit to a linear model to obtain parameters $a$ and $b$, yielding $a = 0.057$ and $b = 0.0064$. **C** Using the relationship obtained in **B**, the time evolution of the coefficient of variation ($CV_\kappa$) during nutrient upshift (starting at $t = 0$) was accurately captured by simulating growth rate and standard deviation.

Growth rate noise was implemented the same way as in the single cell simulations, with an offset value drawn for each cell at birth.

Using the simulation method detailed above, we tracked the number of cells over time, $P(t)$. To calculate the population growth rate, we consider population growth an exponential process and solve for the instantaneous growth rate, $\kappa_{\text{pop}}$. We obtained the instantaneous growth rate by computing the discrete derivative of the natural logarithm of the total number of cells between each time point, specifically

$$\kappa_{\text{pop}} = \frac{\ln(P(t+\Delta t)) - \ln(P(t))}{\Delta t \ln 2} . \tag{41}$$

### V. ESTIMATING DIVISION PROTEIN ALLOCATION FRACTION FROM PROTEOMICS DATA

In order to obtain a value for allocation fraction $f_X$, the identity of the specific cell division proteins, collectively referred to as X proteins, must be known. Although multiple proteins may be involved in setting division timing, experimental evidence suggests that FtsZ is the main determinant of cell division control in *E. coli* [9, 10]. As such, in order to obtain approximate values for $f_X$ we assume that X protein abundance is made up entirely of FtsZ proteins. Obtaining an estimate for $f_X$ requires that the parameters $\alpha$ and $\beta$ be known, given that $f_X = \alpha(\phi_R^{\max} - f_R) + \beta$. In order to estimate these parameters, Eq. (28) can be fit to experimental data, where there are three fitting parameters, $\gamma\alpha$, $\gamma\beta$, and $\kappa_t$ ($\Delta\phi$ can be inferred from experimental data [2]), where $\gamma = \rho_c / X_0 m_X$. To find $\alpha$ and $\beta$ explicitly, the value of $\gamma$ must be known. To estimate its value, we assume that the division threshold, $X_0$, is determined entirely by FtsZ, such that $X_0 = M_X^0 / m_X$, where $M_X^0$ is the total mass of FtsZ at division, and $m_X$ is the mass of a single FtsZ protein. As a result, we

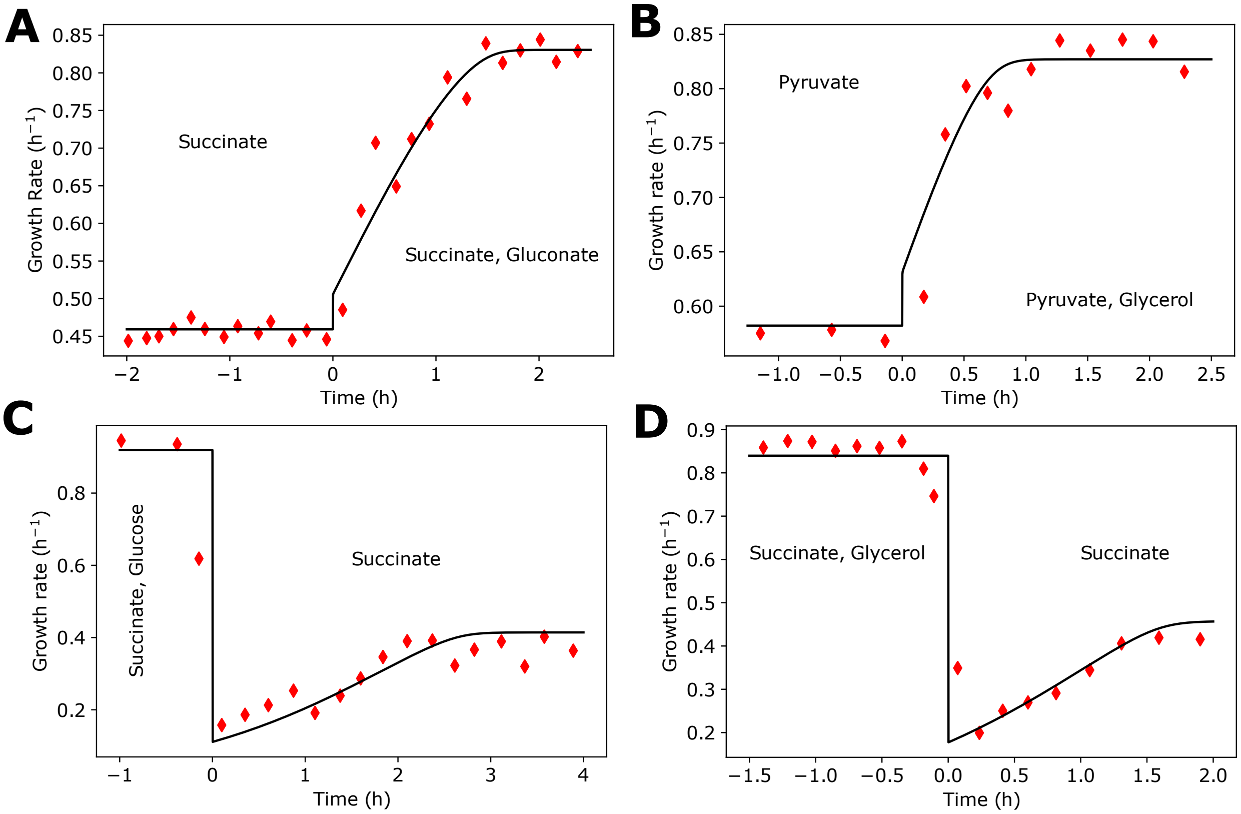


**Supplementary Figure** 3. **Validation of growth rate control model in multiple experimental conditions**. Model successfully predicts growth rate dynamics during nutrient upshift **A,B** and downshift **C,D** in multiple different experimental conditions. Model parameters $\kappa^0_{\mathrm{n,low}}$, $\kappa^0_{\mathrm{n,high}}$, and $\kappa^0_t$ obtained by fitting to data [13] from each experiment.

obtain the expression $\gamma = \rho_c / M^0_X$. Using proteomics data [11], we estimate that $M^0_X \approx 6 * 10^{-4}$ pg. With the typical protein mass density of an *E. coli* cell given by $\rho_c \approx 0.15$ pg/$\mu$m$^3$ [12], the parameters $\alpha$ and $\beta$ can be calculated using $\gamma \approx 250\ \mu\mathrm{m}^{-3}$.

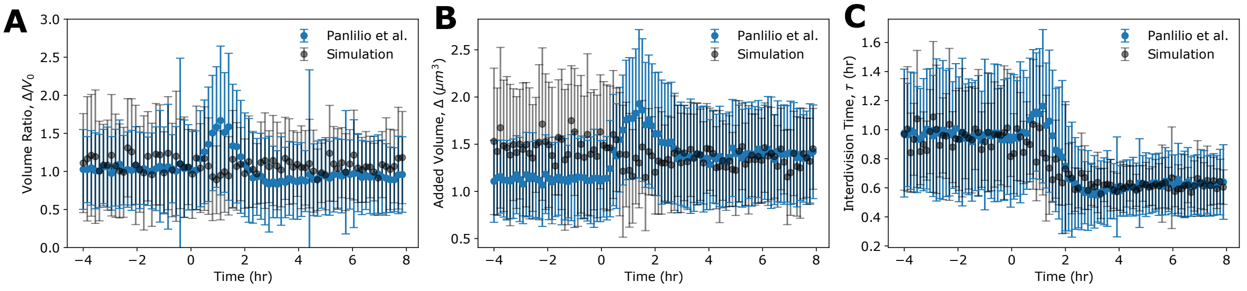


**Supplementary Figure** 4. **Constant $f_X$ model is not able to reproduce experimental results**. **A-C** Generation-averaged dynamics of cell volume ratio (**A**), added volume (**B**), and interdivision time (**C**) from single-cell volume trajectories experiencing nutrient uphshift at $t = 0$ do not agree with experimental data. The constant $f_X$ model predicts that cell size is invariant to nutrient perturbations, thus added volume and volume ratio remain unchanged through nutrient upshift and there is no overshoot in interdivision time. Error bars indicate the standard deviation of the time-binned mean for all time series. Parameters are identical to those used in Figure 3 of the main text (given in Table I), except $\gamma f_X$ remains constant at 0.86 $\mu m^{-3}$.

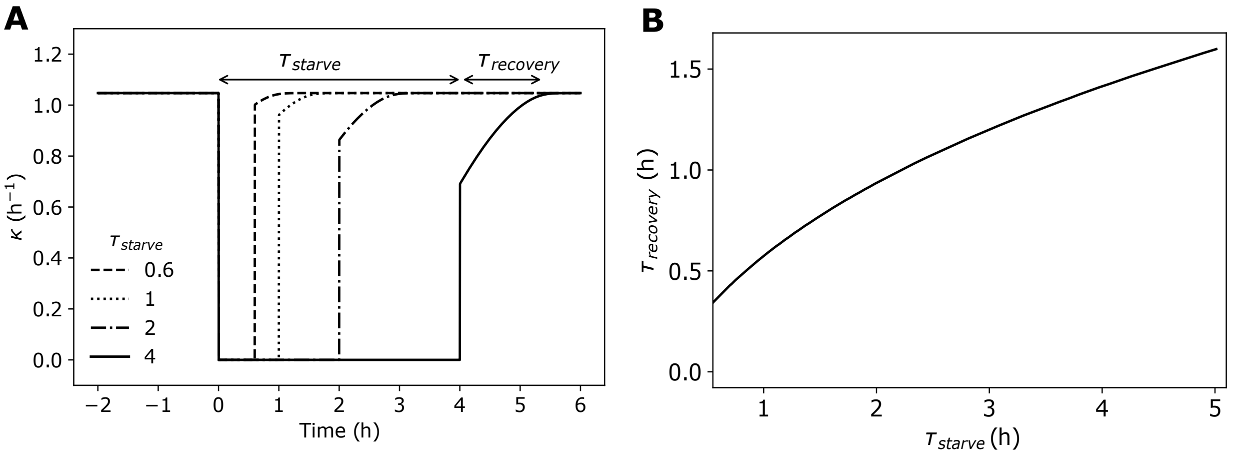


**Supplementary Figure** 5. **Recovery from stationary phase is dependent on starvation length. A** Average single-cell growth rate simulations of bacteria experiencing a nutrient-free pulse of duration $\tau_{\text{starve}}$. For each trajectory with pulse-length $\tau_{\text{starve}}$, the time required following downshift for the growth rate to return to within 99% of the pre-shift level was measured, given by $\tau_{\text{recovery}}$. **B** Quantification of the relationship of $\tau_{\text{starve}}$ and $\tau_{\text{recovery}}$ from the simulations in **A** for two different degradation rates. Parameters are identical to those used in Figure 5 of the main text (given in Table I), except $\kappa^0_{n,high} = 30$ h$^{-1}$.

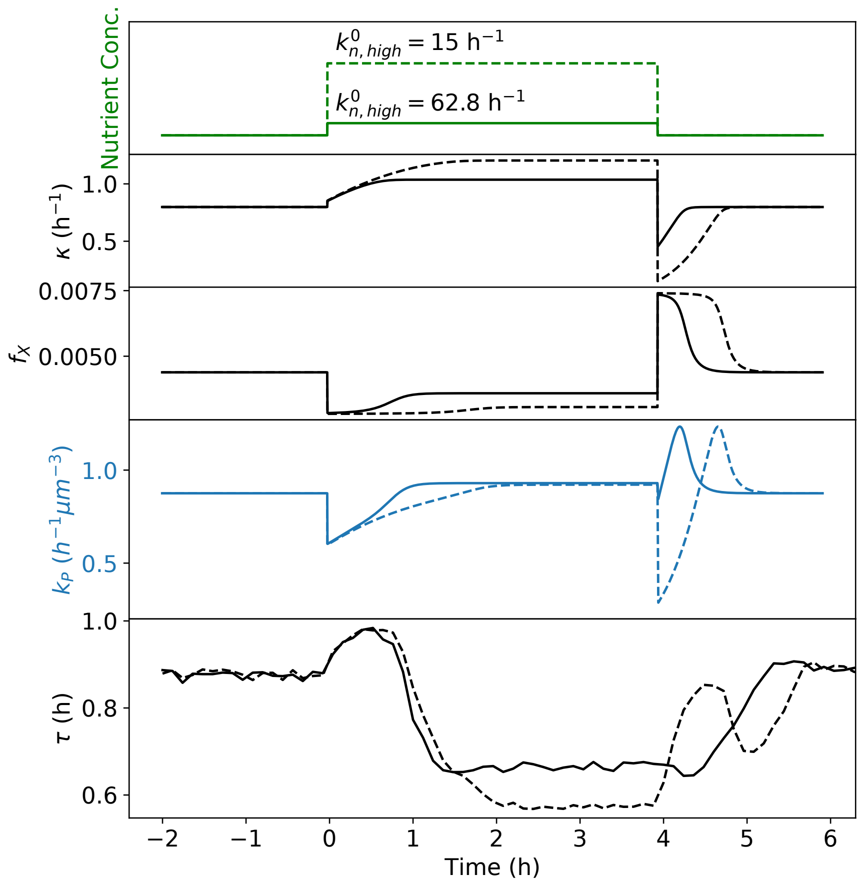


**Supplementary Figure** 6. **Interdivision time recovery behavior is pulse height dependent.** Average single-cell growth rate simulations of bacteria experiencing a nutrient-rich pulse of duration $\tau_{\text{feast}} = 4$, with different nutrient concentrations during the nutrient-rich pulse. In both cases, the top four panels are deterministic simulations of average intracellular dynamics, whereas the bottom panel is the average dynamics of 400 single-cell stochastic simulations. Parameters are identical to those used in Figure 4 of the main text (given in Table I), except for $k^0_{n,high}$ (given on top panel).

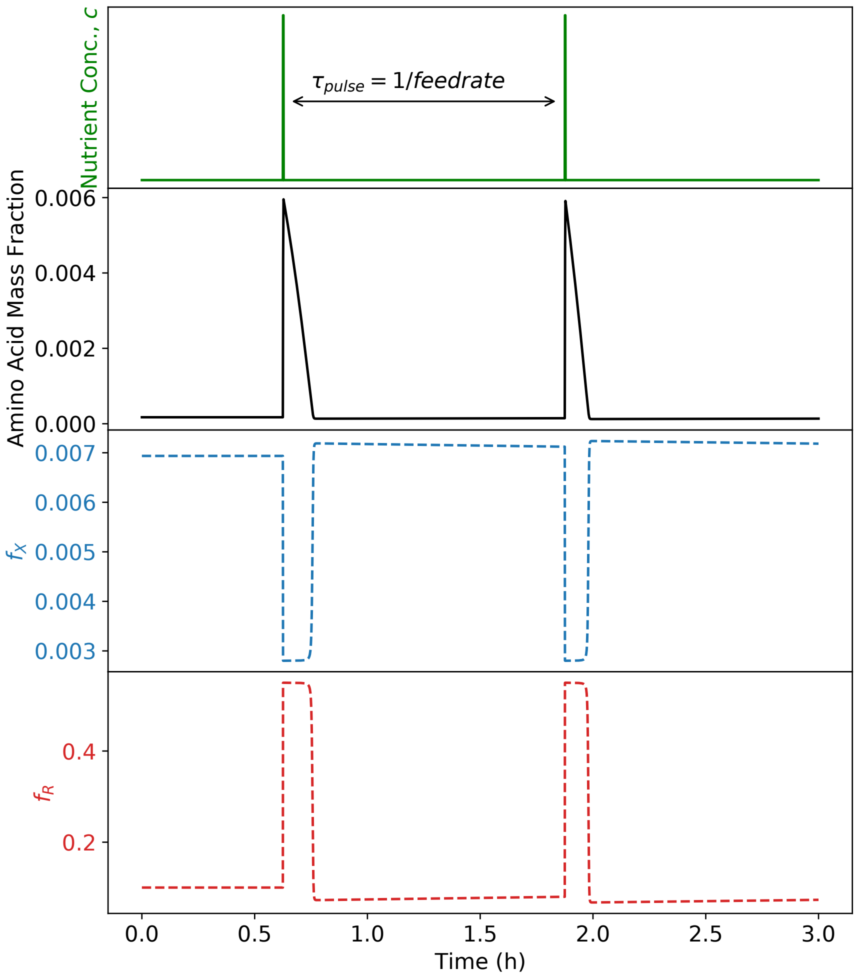


**Supplementary Figure** 7. **Dynamic resource allocation during exit from stationary phase**. Single-cell simulation dynamics of amino acid mass fraction and division protein allocation fraction for *E. coli* experiencing pulses of nutrients with delay $\tau_{\text{pulse}}$ starting from stationary phase. An increase in available nutrients results in an increase in the intracellular amino acid mass fraction. In response to the influx of resources, our model predicts that bacteria transiently prioritize ribosome production ($f_R$) over division ($f_X$) immediately following pulse exposure, similar to nutrient upshift behavior predicted in exponential phase. Parameters are identical to those used in Figure 5 of the main text (given in Table I).

### Supplemental References

[1] N. M. Belliveau, G. Chure, C. L. Hueschen, H. G. Garcia, J. Kondev, D. S. Fisher, J. A. Theriot, and R. Phillips, Cell Systems **12**, 924 (2021).

[2] M. Scott, C. W. Gunderson, E. M. Mateescu, Z. Zhang, and T. Hwa, Science **330**, 1099 (2010).

[3] M. Scott, S. Klumpp, E. M. Mateescu, and T. Hwa, Molecular Systems Biology **10**, 747 (2014).

[4] Y. K. Kohanim, D. Levi, G. Jona, B. D. Towbin, A. Bren, U. A. Correspondence, and U. Alon, Cell Reports **23** (2018), 10.1016/j.celrep.2018.05.007.

[5] P. Wang, L. Robert, J. Pelletier, W. L. Dang, F. Taddei, A. Wright, and S. Jun, Current Biology **20**, 1099 (2010).

[6] M. Osella, E. Nugent, and M. C. Lagomarsino, Proceedings of the National Academy of Sciences of the United States of America **111**, 3431 (2014).

[7] M. Panlilio, J. Grilli, G. Tallarico, I. Iuliani, B. Sclavi, P. Cicuta, and M. C. Lagomarsino, Proceedings of the National Academy of Sciences **118** (2021), 10.1073/PNAS.2016391118.

[8] J. Männik, F. Wu, F. J. Hol, P. Bisicchia, D. J. Sherratt, J. E. Keymer, and C. Dekker, Proceedings of the National Academy of Sciences of the United States of America **109**, 6957 (2012).

[9] K. Sekar, R. Rusconi, J. T. Sauls, T. Fuhrer, E. Noor, J. Nguyen, V. I. Fernandez, M. F. Buffing, M. Berney, S. Jun, R. Stocker, and U. Sauer, Molecular Systems Biology **14**, e8623 (2018).

[10] F. Si, G. Le Treut, J. T. Sauls, S. Vadia, P. A. Levin, and S. Jun, Current Biology **29**, 1760 (2019).

[11] A. Schmidt, K. Kochanowski, S. Vedelaar, E. Ahrné, B. Volkmer, L. Callipo, K. Knoops, M. Bauer, R. Aebersold, and M. Heinemann, Nature Biotechnology **34**, 104 (2016).

[12] R. Phillips, J. Kondev, J. A. Theriot, H. G. Garcia, and N. Orme, *Physical Biology of the Cell (2nd ed.)* (Garland Science, 1998).

[13] D. W. Erickson, S. J. Schink, V. Patsalo, J. R. Williamson, U. Gerland, and T. Hwa, Nature **551**, 119 (2017).
